# Spatial Patterning Analysis of Cellular Ensembles (SPACE) discovers complex spatial organization at the cell and tissue levels

**DOI:** 10.1101/2023.12.08.570837

**Authors:** Edward C. Schrom, Erin F. McCaffrey, Vivek Sreejithkumar, Andrea J. Radtke, Hiroshi Ichise, Armando Arroyo-Mejias, Emily Speranza, Leanne Arakkal, Nishant Thakur, Spencer Grant, Ronald N. Germain

## Abstract

Spatial patterns of cells and other biological elements drive both physiologic and pathologic processes within tissues. While many imaging and transcriptomic methods document tissue organization, discerning these patterns is challenging, especially when they involve multiple elements in complex arrangements. To address this challenge, we present Spatial Patterning Analysis of Cellular Ensembles (SPACE), an R package for analysis of high-plex spatial data. SPACE is compatible with any data collection modality that records values (i.e., categorical cell/structure types or quantitative expression levels) at fixed spatial coordinates (i.e., 2d pixels or 3d voxels). SPACE detects not only broad patterns of co-occurrence but also context-dependent associations, quantitative gradients and orientations, and other organizational complexities. Via a robust information theoretic framework, SPACE explores all possible ensembles of tissue elements – single elements, pairs, triplets, and so on – and ranks the most strongly patterned ensembles. For single images, rankings reflect patterns that differ from random assortment. For sets of images, rankings reflect patterns that differ across sample groups (e.g., genotypes, treatments, timepoints, etc.). Further tools then thoroughly characterize the nature of each pattern for intuitive interpretation. We validate SPACE and demonstrate its advantages using murine lymph node images for which ground truth has been defined. We then use SPACE to detect new patterns across varied datasets, including tumors and tuberculosis granulomas.

## Introduction

Biological tissue function emerges from the local interactions of diverse cells and structures. Whether these interactions occur via direct contact or paracrine signaling, they are determined by the spatial arrangement of the involved elements. Thus, discovering new spatial patterns can reveal new mechanisms of tissue (dys)function [1–5].

Many technologies capture the spatial arrangements of deeply phenotyped cells and structures. Protein-targeted methods, using fluorescence [6–7], sequencing [8], or heavy metal isotopes [9–10], capture 30-100 molecular parameters at submicron resolution. Transcript-targeted methods capture hundreds of molecular parameters at similar resolution [11–13], or the entire transcriptome at lower spatial resolution [14–15]. Multimodal techniques combine the advantages of several methods [16]. In each case, dozens to hundreds of cell types and structures can be identified, in addition to the raw expression of hundreds to thousands of protein or mRNA biomolecules. These spatial elements are too numerous for manual exploration of combinatoric patterns.

Although many computational tools exist to analyze high-plex spatial data, they fall into two broad approaches. The first approach detects quantitative patterns, but only among one or two elements. Marked point processes [17], Gaussian process regression [18–19], and generalized linear spatial models [20] determine whether single biomolecules are spatially variably expressed. Neighbor frequency [21–22], spatial correlation [23–25], and nearest distances [26] determine whether pairs of cell types or biomolecules are significantly co-localized or mutually exclusive. Regardless of their statistical machinery, none of these methods detect patterns involving three or more elements. Moreover, the pairwise methods do not account for non-linear or context-dependent patterns, i.e., patterns that differ qualitatively across tissue regions.

The second approach considers three or more elements simultaneously, but only in terms of discrete microenvironments (MEs) whose internal compositions are assumed homogeneous. Methods of defining MEs are varied, including graph convolutional neural networks [27], hidden Markov random fields [28–29], graph-based community detection [30], the K-means algorithm [31–32] and latent Dirichlet allocation [33–35]. Regardless of the learning algorithm, the discrete nature of MEs cannot capture quantitative features such as smooth gradients or structural orientations.

Thus, despite the sophisticated techniques within each method, the approaches themselves are conceptually limited to specific types of patterns. These “blind spots” overlap: even together, the two approaches do not efficiently detect quantitatively complex or context-dependent arrangements of three or more elements. Moreover, both approaches identify only patterns that differ from null expectations of random assortment within a single specimen, requiring further customized analyses to discern how groups of biological specimens differ systematically from one another.

To overcome these limitations, we present SPACE (Table 1), an information-theoretic algorithm that systematically detects, statistically validates, and thoroughly characterizes spatial patterns of any complexity. SPACE is not merely a performance enhancement within the two existing approaches to incrementally improve the detection of simple patterns. Rather, SPACE is a conceptual invention of a third approach that detects arbitrarily complex patterns, even those in the shared “blind spot” of existing approaches.

**Table 1.**
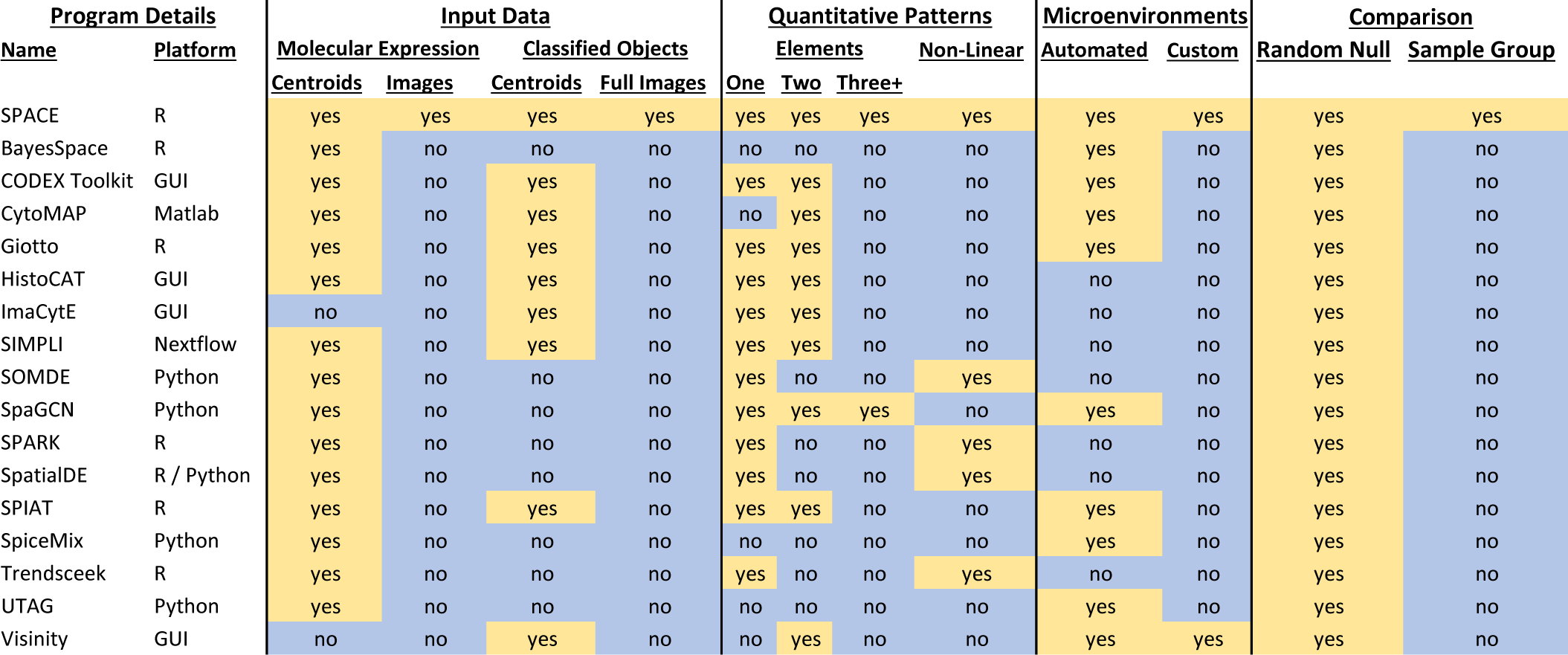
Comparison of SPACE with other spatial analysis platforms. “Input Data” refers to the actual data on which statistical calculations are performed. “Quantitative Patterns” refers to patterns queried directly for single or multiple elements. “Microenvironments” refers to discrete regions with characteristic compositions of elements. “Comparison” refers to whether identified spatial patterns differ from null expectations for a single specimen or differ across sample groups for multiple specimens.

SPACE is compatible with data generated using any collection modality that records values at fixed spatial coordinates, whether 2d pixels, 3d voxels, or tabulated centroids. The values may be categorical objects (e.g., cell types or structures), or quantitative molecular expression levels. For any ensemble of any number of these elements, SPACE quantifies the deviation from random patterning in a single specimen using a metric we call “cis mutual information” (cisMI). Across multiple specimens, SPACE quantifies how an ensemble’s spatial patterning differs across sample groups using a metric we call “trans mutual information” (transMI). SPACE exhaustively explores many ensembles to discern the strongest patterns, which are then characterized in detail to reveal not only co-assortment but also gradients, orientations, and other complex features that may be context-dependent. SPACE is available as an open-source R package.

We demonstrate and validate SPACE on data from different collection modalities, host species and tissues, and input data formats. We focus on highly multiplex imaging of a mouse lymph node where ground-truth spatial patterning is known, as well as 3d volumetric imaging of a mouse tumor, a spatial transcriptome of human intestinal cancer, and a collection of human tuberculosis (TB) granulomas. In addition to rediscovering known patterns, SPACE provides a new level of exploratory power that translates the enormous content of high-plex spatial data into a small set of candidate spatial patterns most likely to reveal important underlying biological processes.

## Results

### SPACE Algorithm

Input data for SPACE can include images of molecular expression, color-coded segmented cells, color-coded pixel-based structures, and/or tables reporting the centroid, categorical type, and average molecular expression of single cells (Fig. 1A). Each input may be 2d or 3d. Multiple inputs representing the same specimen can be analyzed simultaneously to integrate data at the molecular, cellular, and structural levels. Biological elements at any of these levels are simply called “variables.” Across each layer of input data, many 3d spherical or 2d circular neighborhoods are drawn (Fig 1B), and the amount of each variable in each neighborhood is reported in a census (Fig 1C-D). A census may include a variety of neighborhood sizes, for comparison across scales (e.g., adjacent cell contacts at 5-10μm vs. microanatomical zonation at 50-100μm). Contiguous patches of each variable are also recorded and reassorted into hypothetical neighborhoods to create randomized censuses (Fig. 1C), which preserve cellular/structural size and morphology while eliminating spatial patterning (Fig. 1D).

**Figure 1.**
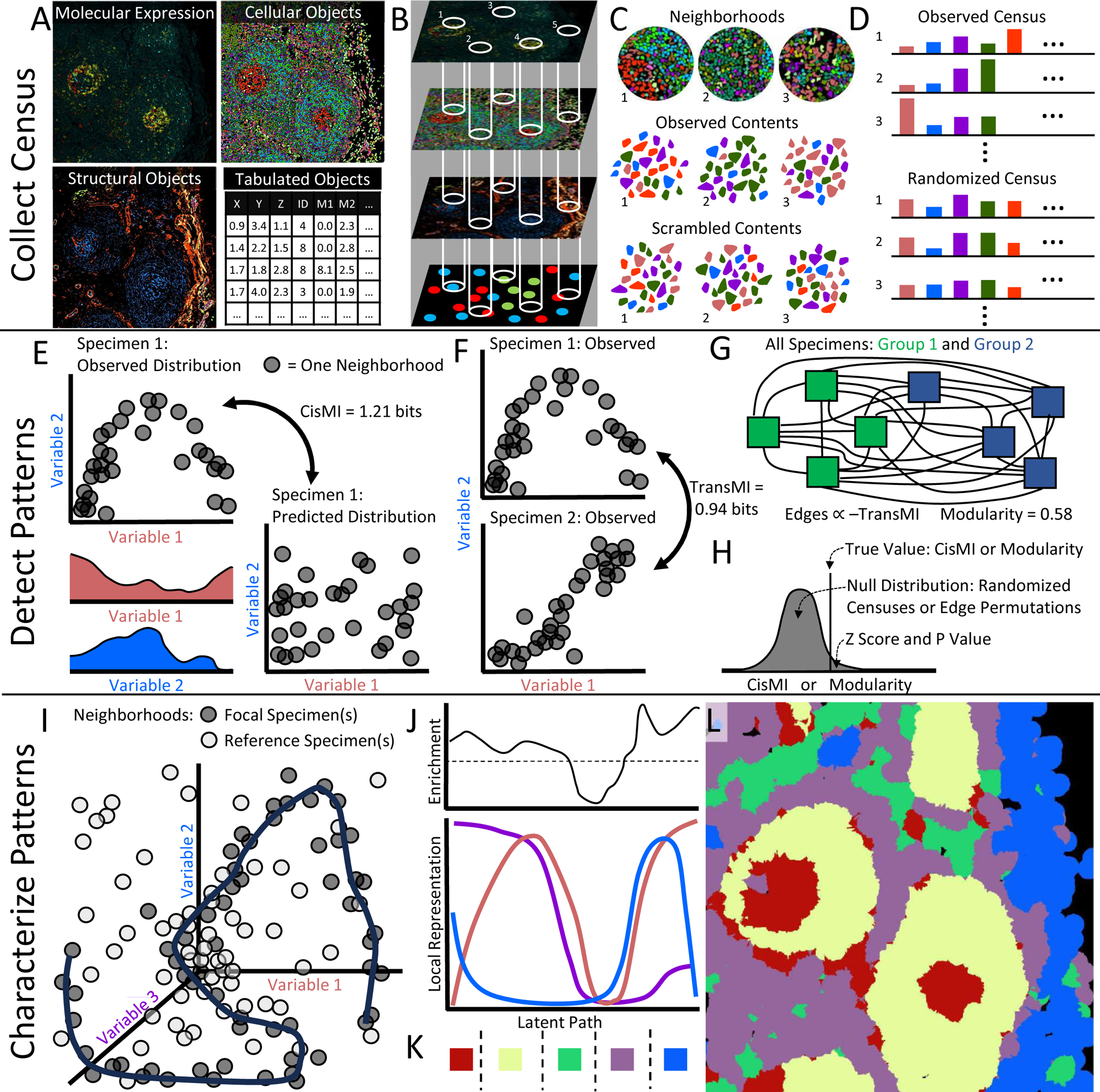
SPACE workflow. A) SPACE accepts images of molecular expression, segmented and classified cells, and classified pixels, as well as tables of cellular centroid position, phenotypic identity, and/or mean molecular expression. Input data may be 3d or 2d. B) Neighborhoods are drawn through each layer of provided data. C) Neighborhood contents are tabulated, as are contiguous patches of each variable, which are scrambled to form randomized neighborhoods. D) True neighborhood contents form the observed census, while scrambled neighborhood contents form randomized censuses. E) CisMI for a given ensemble quantifies the difference between the true distribution of co-occurrence and the distribution predicted without any patterning beyond that which is present in subsets of the ensemble. F) TransMI for a given ensemble begins with the KL Divergence between the observed co-occurrence distributions for each pair of samples. G) KL Divergences form the edge weights on a fully connected network of all samples, from which modularity is calculated. H) CisMI or transMI modularity are expressed as Z-scores relative to a null distribution of the same measures computed on randomized censuses for cisMI or on permuted networks for transMI modularity. I) For a given ensemble, a self-organizing map learns the shape of non-random patterning as a curve through the multi-dimensional co-occurrence distribution from focal sample(s). The co-occurrence distribution for reference sample(s) is ignored. J) The multi-dimensional curve is projected onto the covariation plot revealing specific features of the ensemble’s patterning, as well as enrichment of these features in the focal sample(s) compared to the reference sample(s). L) Manually-defined regions are mapped back onto the tissue for further rounds of SPACE analysis.

Censuses are used to discover patterns that differ from null expectations within a single specimen. For a given ensemble of variables, the observed co-occurrence distributions for the full ensemble and for every subset are used to calculate interaction information. Interaction information is a generalization of mutual information that quantifies the patterning in a group of variables which cannot be explained by any subsets of those variables. Thus, interaction information is essentially the difference between the true distribution of co-occurrence for the full ensemble and the predicted distribution based on all subsets (Fig. 1E). Because this quantifies non-random patterning vs. null expectations within a single specimen, we have named this measure cis-mutual information, or cisMI.

Censuses are also used to discover patterns that differ across groups of specimens. For a given ensemble of variables, the observed co-occurrence distributions for the full ensemble are used to calculate the Kullback-Leibler (KL) divergence between pairs of specimens. KL divergence quantifies the difference between two distributions. Because KL divergence is essentially the difference in patterning for the full ensemble between two specimens (Fig. 1F), we call this measure trans-mutual information, or transMI. TransMI between every pair of specimens yields a fully connected network, where vertices represent specimens and edges are weighted by transMI, such that specimens that share more similar patterning are more closely connected (Fig. 1G). When specimens belong to discrete sample groups, the modularity of this network can be calculated, quantifying how well the ensemble’s spatial patterning distinguishes sample groups.

Once cisMI or transMI modularity have been calculated, they are compared against a null distribution (Fig. 1H). For cisMI, the null distribution is constructed by repeating the cisMI measurement on at least 100 randomized censuses, to control for apparent patterning that is really a byproduct of tissue geometry or compositional abundances. For transMI modularity, the null distribution is constructed by permuting the network edge weights at least 100 times, to control for apparent modularity that is really a byproduct of outlier specimens or small samples sizes. In either case, the original cisMI or transMI modularity compared against the null distribution yields a Z score and P value. These calculations are repeated for many ensembles to yield many P values, which are ranked to reveal ensembles with the most significant patterning.

Low P values and high cisMI or transMI modularity identify ensembles of interest, but they do not describe the ensemble’s patterning. To characterize a significant ensemble, the distribution of co-occurrence is generated, for both focal and reference censuses. For cisMI, the observed census is focal and a randomized census is the reference. For transMI, the pooled censuses from one sample group are focal, and pooled censuses from another sample group are the reference. A self-organizing map places a curve through the areas of highest focal neighborhood density (Fig. 1I). The curve is projected into lower dimension by plotting the value of each variable along the curve separately. The resulting covariation plot shows how each variable in the ensemble covaries as the tissue is traversed along a hidden but illustrative (i.e., latent) path (Fig. 1J). This reveals context-dependent associations, quantitative gradients, and other features of an ensemble’s spatial patterning in the focal specimen(s). The density of focal neighborhoods along the latent path is compared to the density of reference neighborhoods, to show which features of the ensemble’s spatial pattern are specifically enriched or depleted in the focal specimen(s) vs. the reference (Fig. 1J).

To visualize features of the ensemble’s spatial pattern in the original tissue context, manual breakpoints can be drawn on the covariation plot, splitting the image into MEs with different amounts or gradients of the involved variables (Fig 1K). These MEs are mapped back onto the original image with new coloring (Fig 1L). Because MEs are formatted identically to other SPACE inputs, they can be recycled for further analysis of their own spatial patterning.

### SPACE Detects Known Patterns and Gradients in the Mouse Lymph Node

To compare SPACE output against ‘ground-truth’ for a well-studied tissue, we used a 42-plex image of a mouse popliteal lymph node (LN) captured with the IBEX protocol [36] (Fig. S1A). Using CellPose 2.0 [37], we segmented individual cells (Fig. S1B). To restrict our focus to well-studied cellular subsets, we chose 20 lineage markers to define canonical expression profiles (Table S1) and deployed a semi-supervised pipeline (Fig. S1C-H) to classify the cells into 19 types (Fig. 2A) and map them onto a segmentation mask (Fig. 2B). On this mask, we censused neighborhoods of radius 10 µm to capture close cellular contacts. After examining every ensemble of one, two, or three cell types, there were 24 ensembles with patterning that was both substantial (cisMI > 0.05 bits) and significant (P < 0.05 after Benjamini-Hochberg correction for multiple testing) (Fig. 2C). Performing the same analysis on a table of cell centroids (Table S2) yields similar results (Fig. S2).

**Figure 2.**
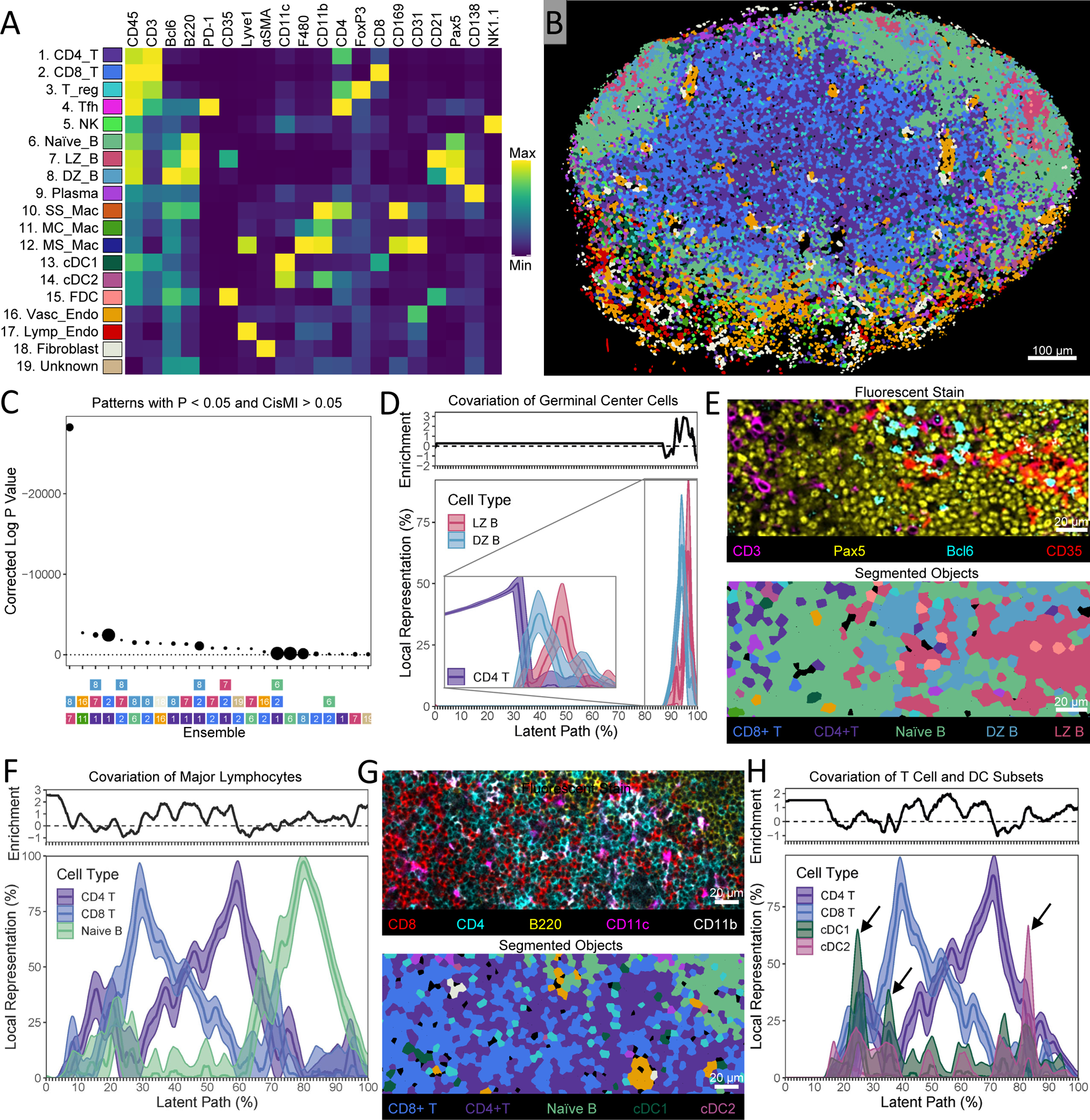
Demonstration of SPACE on a mouse LN. A) Marker expression profiles of 19 segmented cell types. B) Segmentation image used in SPACE analysis. C) Ranking of the most significant non-randomly patterned ensembles by corrected P-value, where point size is proportional to cisMI magnitude in bits. D) Covariation plot for LZ and DZ B cells, including smoothed mean and smoothed 95% confidence interval as well as continuous enrichment score vs. randomized expectations. The same covariation plot with the addition of CD4 T cells is shown in the inset. E) Visual example of the orientation of GCs with the DZ facing toward the T zone and the LZ facing away from it. Fluorescent stain and segmented objects represent the same tissue area. F) Covariation plot for CD4 T cells, CD8 T cells, and naïve B cells, including smoothed mean and smoothed 95% confidence interval as well as continuous enrichment score vs. randomized expectations. G) Visual example of the opposing gradient of CD4 and CD8 T cells with respect to the naïve B follicle. Fluorescent stain and segmented objects represent the same tissue area. H) Covariation plot for CD4 T cells, CD8 T cells, cDC1s, and cDC2s, including smoothed mean and smoothed 95% confidence interval as well as continuous enrichment score vs. randomized expectations. Black arrows highlight peaks in cDC abundance.

Although not immunized, the mouse was conventionally housed, and so we expected the patterns detected to reflect both the structure of a resting LN as well as features of mild immune activity. Indeed, the most significant ensemble involves germinal center (GC) B cells from both the dark zone (DZ) and light zone (LZ) (Fig. 2C). The covariation plot shows that DZ and LZ B cells are concentrated together in less than 15% of the image, a feature which is highly enriched compared to null expectations (Fig. 2D). Within these concentrated regions, there are areas where DZ B cells dominate in abundance over LZ and vice-versa. Together, these features of the covariation plot represent the GC and its well-known subdivision into the DZ and LZ.

The third most significant ensemble involves CD4 T cells in addition to DZ and LZ B cells, indicating that these two GC B cell subsets share further patterning relative to CD4 T cells. (Note that these are paracortical CD4 T cells, not follicular helper T cells, which were too rare to create statistically significant patterning.) A second covariation plot including CD4 T cells reveals that within the GC, DZ B cells tend to be proximal to the highest concentrations of CD4 T cells (in the paracortical T zone), while LZ B cells tend to be distal (Fig. 2D, inset). Indeed, the orientation of GCs with the DZ facing the T zone can be seen in this image (Fig. 2E). While the coherence and internal structure of the GC can be explained by DZ and LZ B cells only, significant cisMI for the three-variable pattern is statistical evidence that GC orientation is also non-random.

We also investigated the three-variable pattern with the largest cisMI, involving CD4 T cells, CD8 T cells, and naïve B cells (Fig. 2C). These populations define the largest microanatomical regions of the LN – the paracortical T zone and primary B follicles. Indeed, the covariation plot shows separate regions where T cells vs. naïve B cells dominate (Fig. 2F, 15-65% vs. 65-100% along the latent path). Additionally, within the T zone, CD4 and CD8 T cells form opposing gradients: CD4 T cells dominate closer to main B follicle and CD8 T cell dominate further away. This opposing gradient of T cells is one of the most enriched features of the covariation plot compared to null expectations (Fig. 2F), and it can be seen in the image (Fig. 2G). Although the visual manifestation of this pattern depends on the slicing angle of the LN (Fig. S3), it remains that CD4 and CD8 T cells co-occur at large but oppose one another specifically within the B-cell-devoid T zone. This context-dependent gradient of paracortical naïve CD4 and CD8 T cells was discovered recently [38] using manual histo-cytometry analyses [39], and SPACE recovers it via automated exploration.

Additionally, Type 1 dendritic cells (cDC1s) peak in abundance closer to the center of the T zone where CD8 T cells are concentrated, whereas Type 2 dendritic cells (cDC2s) peak in abundance at the edge where CD4 T cells are concentrated [38–39]. This is a four-element pattern, which we queried directly in SPACE. Indeed, CD4 T cells, CD8 T cells, cDC1s, and cDC2s together have significant cisMI (P = 0.023). The covariation plot reveals that cDC1s peak in abundance in the part of the T zone where CD8 T cells dominate, whereas cDC2s peak in abundance where CD4 T cells dominate (Fig. 2H).

Quantitative Multi-Variable Patterns Are Inaccessible to Existing Analysis Approaches Existing spatial analysis platforms (Table 1) fall into two broad categories: the quantitation of homotypic or pairwise heterotypic patterns, or the delineation of categorical MEs. While both approaches can provide useful insights [17–35], we examined whether they could detect the same patterns as SPACE. We applied examples of these two categorical approaches to the mouse LN to search for the GC orientation and T cell and DC gradients detected by SPACE. Although many statistical procedures exist for pairwise analyses, we provide a generalized example that shows why homotypic and pairwise heterotypic patterns are conceptually incapable of capturing the patterns that SPACE finds. Likewise, although many learning algorithms exist to define MEs, one generalized example shows why discrete MEs are conceptually incapable of capturing the patterns that SPACE finds.

Using the table of centroids and phenotypes for segmented cells (Table S2), we censused each cell’s neighbors within 10 μm using the true phenotypes as well as permuted phenotypes to describe null expectations in the absence of spatial patterning (Fig 3A). We then counted how often each pair of cell types were neighbors in reality vs. in the permutations (Fig. 3B). CD4 T cells, CD8 T cells, naïve B cells, LZ B cells, and DZ B cells all have significantly more homotypic neighbors than expected (Fig. 3C, gray boxes), reflecting their assortment into distinct tissue regions. CD4 and CD8 T cells neighbor one another more frequently than expected, and they each neighbor naïve B cells less frequently than expected (Fig. 3C, white boxes). This reflects the coexistence of CD4 and CD8 T cells in the T zone separate from B follicles. However, it fails to suggest that CD4 and CD8 T cells form opposing gradients within the T zone, because these opposing gradients are context-specific to the T zone and contradict the tissue-wide co-occurrence pattern. Similarly, both LZ and DZ GC B cells neighbor CD4 T cells less frequently than expected (Fig. 3C, black boxes), reflecting the separation of GCs from the T zone. However, there is no indication that DZs preferentially face toward the T zone compared to LZs, because this requires information on all three of the involved cell types simultaneously. Three-variable patterns often cannot be derived from purely pairwise measurements.

**Figure 3.**
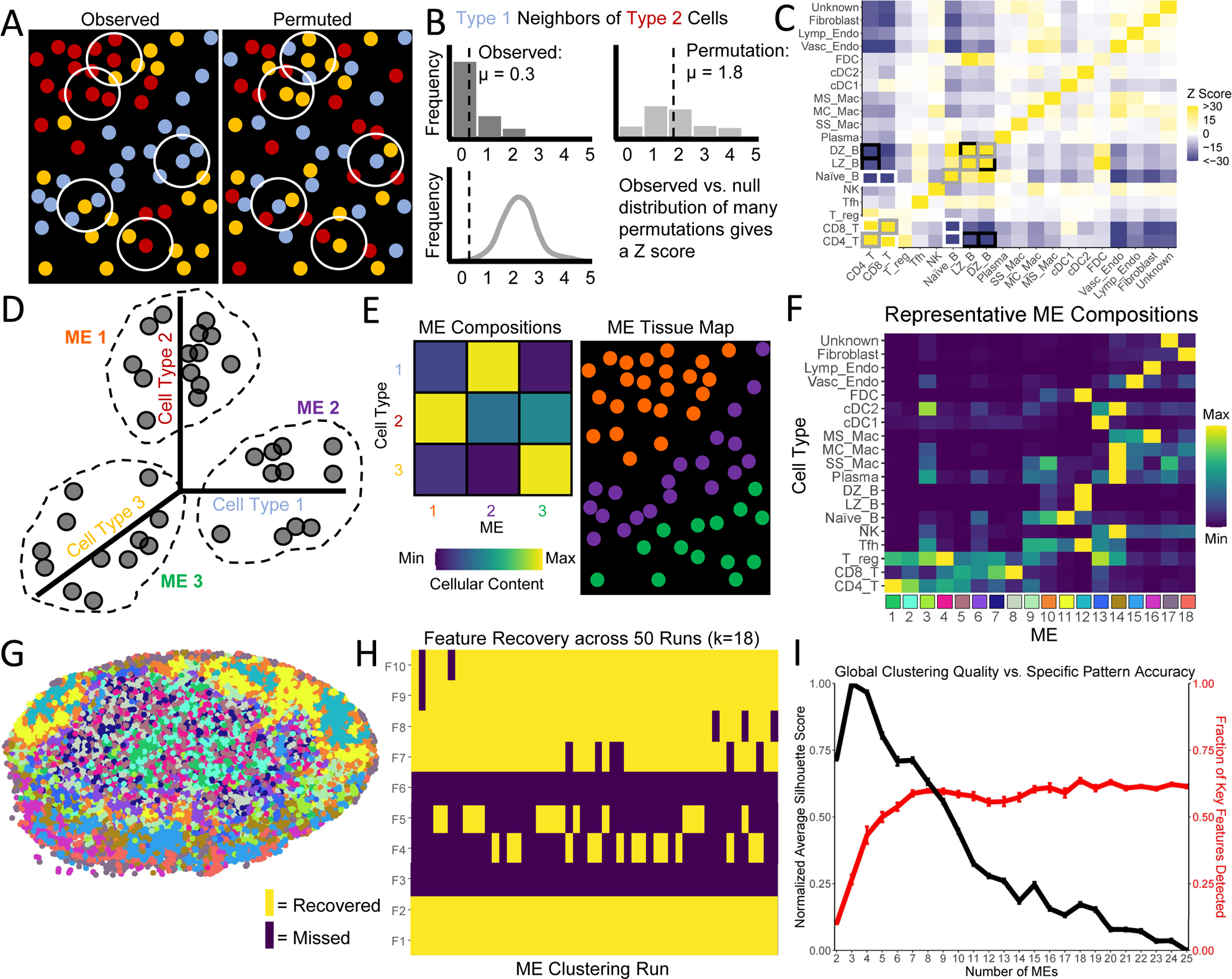
Application of existing spatial analysis techniques to the mouse LN. A) Cartoon of observed and randomized census collection from 2d centroid tables. B) Cartoon of pairwise analysis comparing neighbor frequency to null expectations. C) Pairwise analysis of the neighbor frequency in the mouse LN. Z-scores are relative to 100 random permutations of cellular phenotypes. Gray boxes highlight self-clustering of T cells and B cells. Black boxes highlight co-occurrence of DZ and LZ GC B cells and their separation from T cells. White boxes highlight co-occurrence of CD4 and CD8 T cells and their separation from naïve B cells. D) Cartoon of ME definition via clustering-based machine learning. E) Cartoon of ME composition and visual mapping. F) ME compositions from a representative k-means clustering seeking K=18 MEs. G) Visual mapping of the representative clustering. H) Key feature recovery for each of 50 clustering runs with K=18. I) Mean silhouette scores for the quality of clustering (black) and mean fraction of key features recovered (red) for repeated clustering runs across a range of K values. Error bars represent standard error.

We also used machine learning to define MEs of consistent cell type compositions (Fig. 3D). Although we use the K-means algorithm here, any learning algorithm yields similar information: the mean cellular composition of each ME and a visual map of which cells belong to each ME (Fig. 3E). For a representative clustering with K=18 MEs, the cellular compositions (Fig. 3f) and the visual map (Fig. 3G) are shown. It is conceptually obvious that discrete MEs cannot capture smooth cellular gradients in a strict sense. Nonetheless, we translated the gradients and orientations identified by SPACE into 10 key features that could be suggested by discrete MEs in principle (Table S3). For example, if the ME(s) enriched for CD4 T cells are also enriched for CD8 T cells, then they likely represent the T cell zone. Among these MEs, if CD4 T cell content is negatively correlated with CD8 T cell content, one might argue that the MEs demonstrate opposing gradients (Fig. 3F, MEs 1-8). But other key features may be missing; for example, if LZ GC B cells and DZ GC B cells are most enriched in the same ME (Fig. 3F, ME 12), then the MEs do not distinguish LZ vs. DZ and therefore cannot possibly indicate GC orientation. Across 50 K-means clustering runs with K=18, we found that different runs capture different key features, but no single run captures more than 7 of the 10 key features (Fig. 3H).

A priori, there is no reason to seek exactly 18 MEs, so we varied K from 2 to 25 and performed 50 runs at each value. The average silhouette score, which measures global clustering quality to guide the choice of K, peaks at 3 (Fig. 3I, black line). However, when K=3, only 26.6% of key features are recovered on average. Recovery of key features peaks at K=18, but still only 63.6% of these features are recovered on average (Fig. 3I, red line), and the silhouette score is suboptimal. Different key features tend to be recovered at different K values (Fig. S4A), but no single clustering run at any K value recovered more than 8 of 10 (Fig. S4B).

It is possible that a rare K-means run at K=18 might have recovered all 10 key features. Indeed, a more sophisticated algorithm might routinely recover all 10 key features. This possibility does not mean that MEs capture the complex quantitative patterns that SPACE detects, for two reasons. First, the 10 key features used here derive from only three of 644 significant patterns included in the full exploratory output of SPACE (only 24 with the highest cisMI magnitude are shown in Fig. 2C) – many more key features could be defined to represent well-determined patterns in the mouse LN. It is unlikely that a reasonable number of MEs could simultaneously encode evidence of many more patterns, no less all 644, particularly the more subtle patterns with small but non-zero cisMI. Second, the 10 key features were only identified from the ME compositions because they were expected in advance. In an exploratory setting, SPACE would still report all these significant patterns, but it would be unclear which features of ME compositions warrant serious biological interpretation. For example, if two MEs were enriched for both LZ and DZ GC B cells, a slight preponderance of DZ GC B cells in the ME with the higher fraction of CD4 T cells (Key Feature 9) would likely be dismissed as noise, rather than a meaningful indicator of GC orientation. It is only because of the statistically validated SPACE output that we know how to interpret such a small detail in the ME output. While discrete MEs usefully highlight general trends of co-assortment, they do not reliably detect complex quantitative features of patterning, such as the gradients discussed above.

### SPACE Operates on a Three-Dimensional Input Data

To demonstrate SPACE’s compatibility with 3-dimensional spatial data, we used an 808 µm x 753 µm x 270 µm volume of a murine MC38 tumor imaged using the Ce3D protocol [40]. Based on fluorescent markers of CD3, CD4, CD8, and CD31, we generated a 3d rendering of CD4 T cells, CD8 T cells, and blood vessels in the tumor (Fig. 4A). After censusing at a length scale of 50 µm to capture broad patterns of cellular association, SPACE detects a significant pattern involving all three variables (cisMI = −0.17 bits, P = 1.01×10^−108^). We investigated this pattern further with a covariation plot (Fig. 4B).

**Figure 4.**
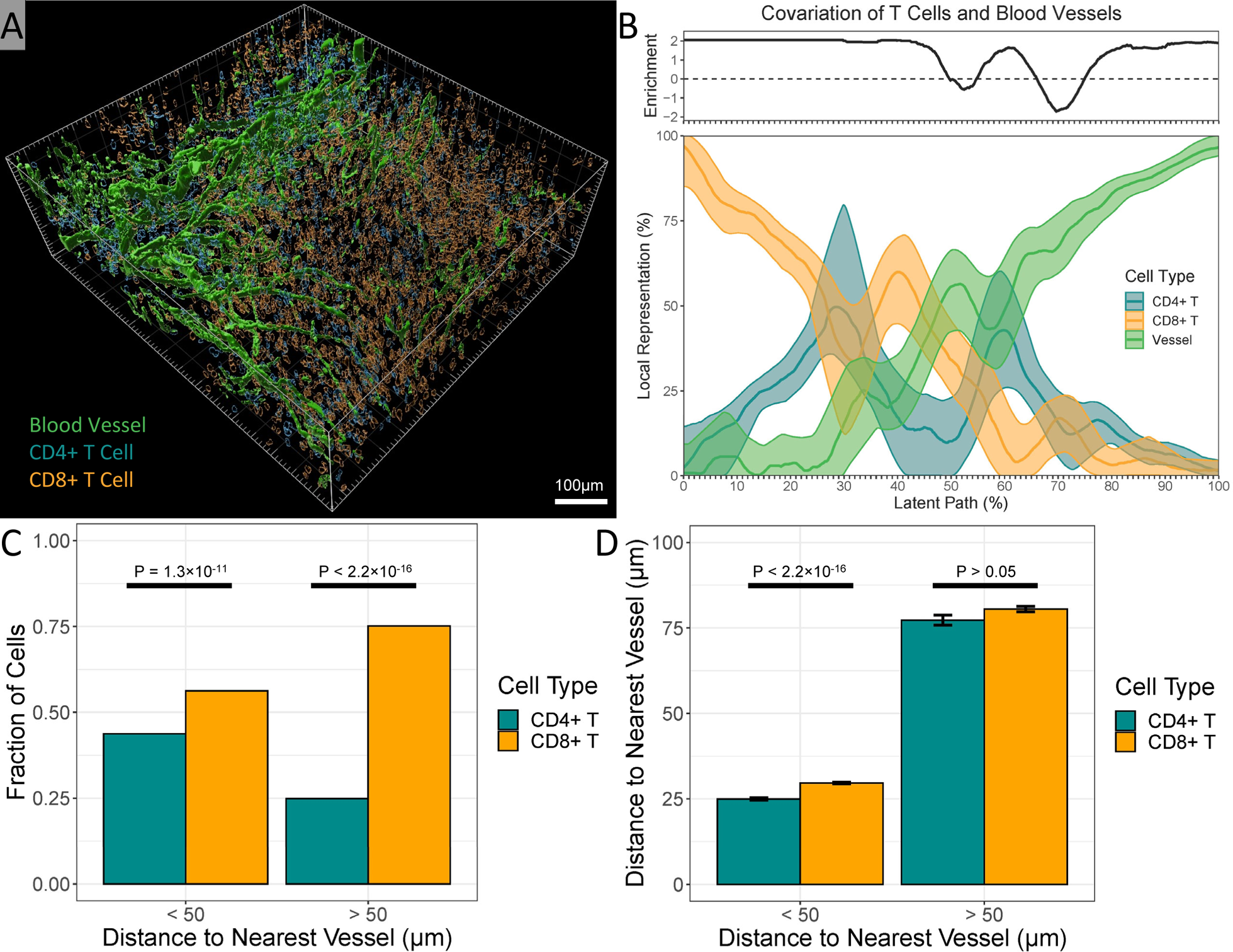
Demonstration of SPACE on a 3d mouse tumor volume. A) Three-dimensional rendering of the tumor volume. B) Covariation plot for CD4 T cells, CD8 T cells, and blood vessels, including smoothed mean and smoothed 95% confidence interval as well as continuous enrichment score vs. randomized expectations. C) Fraction of T cells that are CD4 vs. CD8, separated into cells < vs. > 50 µm from the nearest blood vessel, compared using a binomial linear model. D) Distance to the nearest blood vessel compared between CD4 and CD8 T cells, separated into cells < vs. > 50 µm from the nearest blood vessel, compared using a Gaussian linear model.

Many features of the covariation plot are informative. Much of the tumor volume is dominated either by T cells only (mostly CD8 T cells; 0-25% of latent path) or by blood vessels only (75-100% of latent path). Both features are heavily enriched compared to null expectations. This indicates not only the obvious fact that T cells and blood vessels are separate structures, but also that CD8 T cells outnumber CD4 T cells, especially in regions devoid of blood vessels, which we confirm via direct measurement (Fig. 4C).

Regions where blood vessel content is intermediate (25-75% of latent path) reveal more subtle patterns. Where vessel content is medium-low, CD8 T cells equal or outnumber CD4 T cells (25-50%), and this is enriched vs. null expectations. Meanwhile, where vessel content is medium-high (50-75%), regions where CD4 T cells outnumber CD8+ T cells are also enriched (55-65%), but regions where CD8 T cells equal CD4 T cells (50-55%, 65-75%) are depleted. This indicates that specifically in the regions bordering blood vessels, CD4 T cells preferentially sit closer to vessels, which we confirm via direct measurement (Fig. 4D). While the confirmations (Fig. 4C,D) require custom analysis and code, all these nuanced details are available from a single SPACE command. Recent 3D intravital imaging suggests that peri-vascular collections of different T cell subsets arise in tumors [41] and after vaccination [42], consistent with SPACE’s findings.

### SPACE Operates on Spatial Transcriptomic Input Data

To demonstrate SPACE’s compatibility with spatial transcriptomic data, we used results from a 6.5mm x 6.5mm section of human intestinal colorectal cancer obtained using the Visium Spatial Gene Expression protocol, publicly available from 10x Genomics. Filtering for transcripts that appear in at least half of the 2660 spots yielded 6,914 transcripts. After censusing at a length scale of 300 µm to capture large groups of neighboring spots, we used SPACE to quantify cisMI for each individual transcript. CisMI correlated very strongly with the global Moran’s I for each transcript published by 10x Genomics (Fig. 5A).

**Figure 5.**
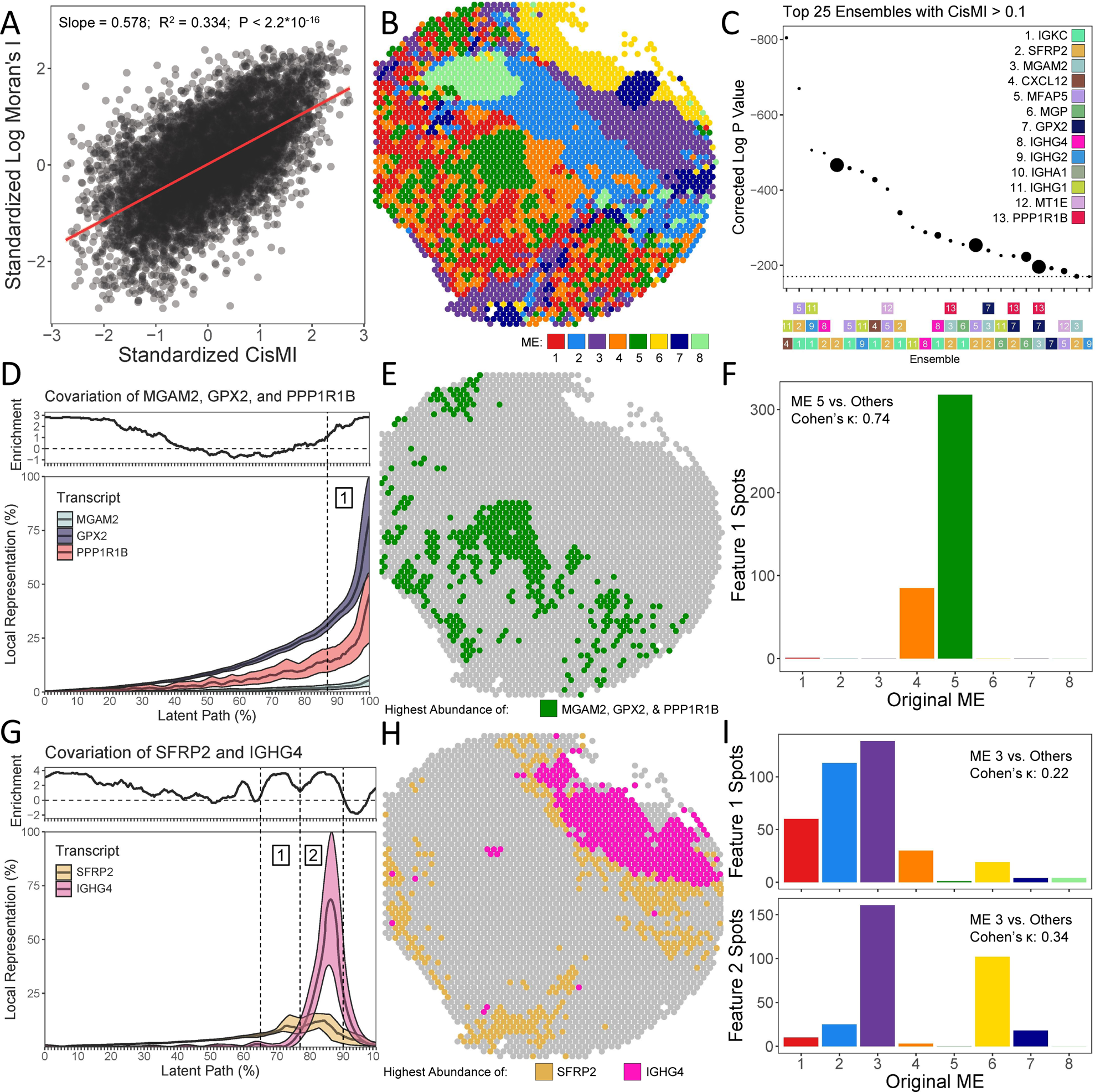
Demonstration of SPACE on spatial transcriptomic spots from human intestinal cancer. A) Comparison of Moran’s I vs. cisMI for 6878 single transcripts, compared using a Gaussian linear model. B) Categorization of 2660 spatial spots into 8 MEs via k-means clustering. C) Ranking the most significant non-randomly patterned ensembles of differentially abundant transcripts by corrected P-value, where point size is proportional to cisMI magnitude in bits. D) Covariation plot for MGAM2, GPX2, and PPP1R1B, including smoothed mean and smoothed 95% confidence intervals as well as continuous enrichment score vs. randomized expectations. E) Mapping of the covariation feature at 87-100% of the latent path onto the original 2660 spatial spots. F) Fraction of spots in the covariation feature that belong to each of the original MEs. G) Covariation plot for SFRP2 and IGHG4, including smoothed mean and smoothed 95% confidence intervals as well as continuous enrichment score vs. randomized expectations. H) Mapping of the covariation features at 65-77% and 77-90% of the latent path onto the original 2660 spatial spots. I) Fraction of spots in each covariation feature that belong to each of the original MEs.

In addition to quantifying the patterning of each transcript individually, 10x Genomics also used k-means clustering to group the spots into 8 different MEs (Fig. 5B). A total of 13 transcripts are differentially abundant vs. their at-large mean in at least 4 of the 8 MEs. Among these 13 transcripts, we censused at a length scale of 150 µm to capture directly neighboring spots, and we measured cisMI for ensembles up three transcripts (Fig. 5C).

The 3-element ensemble with the highest cisMI score includes the transcripts MGAM2, GPX2, and PPP1R1B. The covariation plot reveals that all three transcripts are positively correlated (Fig. 5D), even beyond their pairwise correlations (Fig. S5A-C). In particular, the tissue region highly enriched for all three transcripts (87-100% along the latent path) corresponds closely to ME 5 (Fig. 5E). The gated region on the covariation plot and ME 5 share a Cohen’s κ of 0.74 [0.70, 0.77], indicating “good” to “excellent” agreement (Fig. 5F) [43].

In addition to recovering spatial patterns that are accessible to existing methods, SPACE also finds new patterns. For example, one of the top 5 significant ensembles includes the transcripts SFRP2 and IGHG4. The covariation plot reveals that these transcripts are positively correlated except at peak abundance, where instead there are two separate and highly enriched regions: one where SFRP2 dominates over IGHG4 and one where the opposite is true (Fig. 5G). This is not indicated by the overall positive correlation of SFRP2 and IGHG4 (Fig. S5D). Mapping these two regions back onto the tissue reveals that neither region corresponds well with any of the MEs (Fig. 5H). Although both regions correspond best with ME 3, the gated regions on the covariation plot and ME 3 share Cohen’s κs of only 0.22 [0.18, 0.27] and 0.34 [0.29, 0.39], indicating “poor” agreement [43]. Neither pairwise correlation nor discrete MEs capture this highly significant context-dependent pattern between SFRP2 and IGHG4, but SPACE does. Because the downregulation of SFRP2 is a marker of colorectal cancer [44–45], this pattern agrees with known role of immunoglobulin-expressing cells in infiltrating and combating the tumor [46–47].

### SPACE Directly Compares Spatial Patterning across Sample Groups

Ultimately, the most important spatial patterns may be those that differ systematically among groups of specimens. To address this need, SPACE computes not only cisMI to detect patterning within a single specimen but also transMI to measure detect patterning that distinguishes speciments from different sources, treatment conditions, patient states, and so on. To demonstrate the utility of transMI, we analyzed a publicly available data set of human TB granulomas. Using a 37-plex MIBI-TOF panel, two 500μm x 500μm fields of view (FoVs) were imaged from each of 15 human TB patients [34]. Of these patients, nine were undergoing diagnostic biopsies, three were undergoing therapeutic resections, and three were sampled postmortem. This range of clinical statuses may represent progressing disease. The original study uncovered patterns of granuloma organization that transcend clinical status (e.g. eight MEs with unique cell type compositions). Here, we identify further patterns of granuloma organization that differ across clinical statuses.

Although not many patterns of cellular abundance correlate with clinical status [34], SPACE also measures two metrics of cellular diversity. Alpha diversity, which measures the number and evenness of cell types in each FoV (Fig. 6A), is higher for biopsy and resection samples compared to postmortem material (Fig 6B). Beta diversity, which measures the distinctiveness ME cellular compositions (Fig. 6C), is higher in biopsy samples, compared to either resection or post-mortem (Fig. 6D).

**Figure 6.**
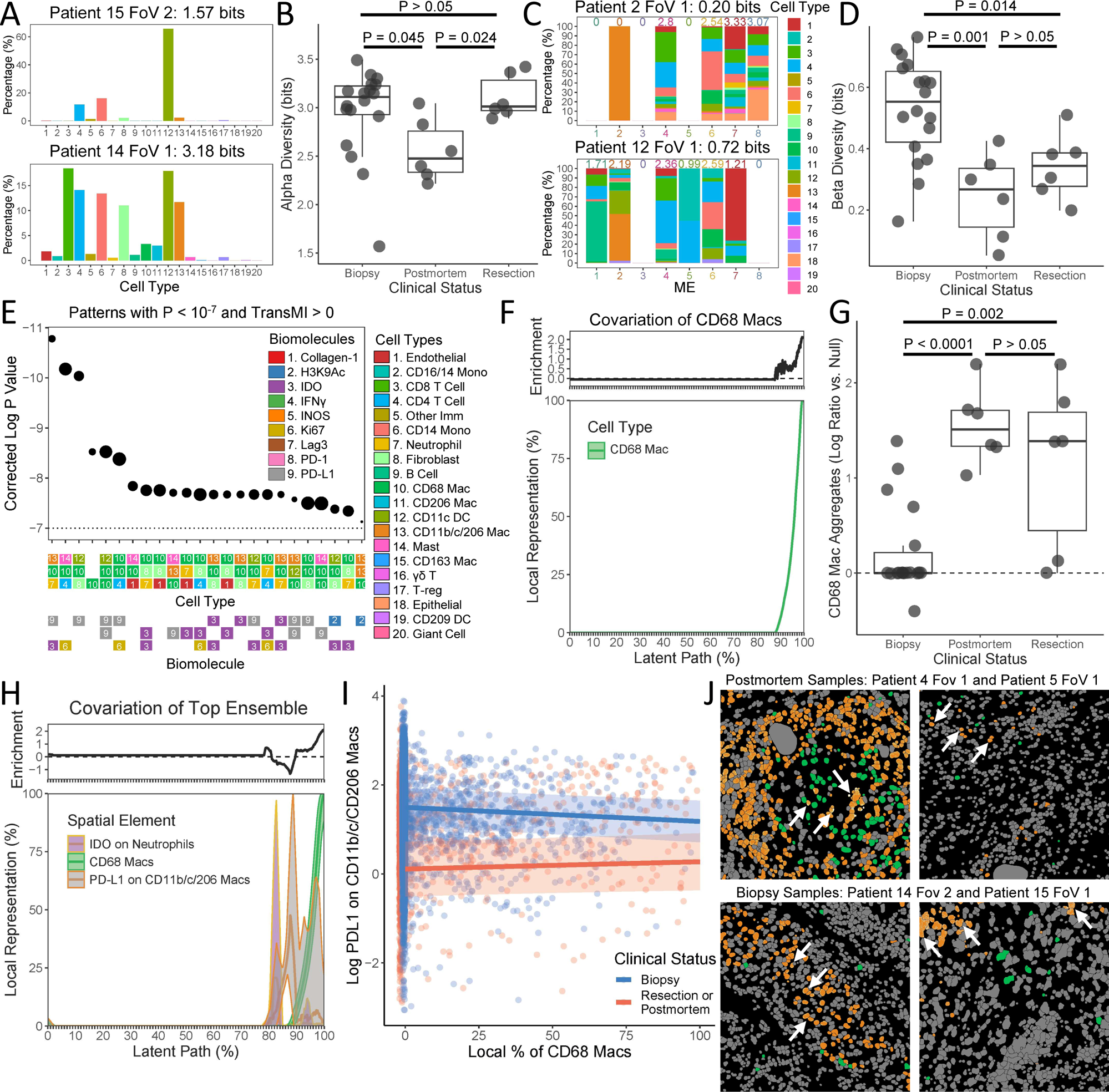
Demonstration of SPACE on a 15-patient data set of human TB granulomas. A) Example FoVs with low and high cellular alpha diversity. B) Comparison of cellular alpha diversity across clinical statuses, compared using a Gaussian linear model. C) Example FoVs with low and high cellular beta diversity across MEs. D) Comparison of cellular beta diversity across clinical statuses, compared using a Gaussian linear model. E) Ranking of the ensembles most differently patterned across clinical statuses by corrected P-value, where point size is proportional to transMI modularity. F) Covariation plot for CD68 macs alone in postmortem and resection samples, including smoothed mean and smoothed 95% confidence intervals as well as continuous enrichment score vs. biopsy samples. G) Comparison of fraction of neighborhoods containing CD68 mac aggregates relative to null expectations across clinical statuses, compared using a Gaussian linear model. H) Covariation plot of neutrophil-specific IDO expression, CD68 mac abundance, and TP mac-specific PD-L1 expression in postmortem and resection samples, including smoothed mean and smoothed 95% confidence intervals as well as continuous enrichment score vs. biopsy samples. I) Change in TP mac-specific PD-L1 expression as a function of local CD68 mac abundance, separated into biopsy vs. postmortem or resection samples, compared using a Gaussian linear mixed effect model. J) Visual examples of high TP mac-specific PD-L1 expression near CD68 macs in postmortem and resection samples, but away from CD68 macs in biopsy samples. White arrows indicate three of the top PD-L1 expressing TP macs in each FoV.

TransMI highlights more specific patterns that differ across clinical statuses. We censused each FoV at a length scale of 10µm to capture close cellular interactions and measured transMI for ensembles of up to three cell types and/or cell-type-specific functional markers. Strikingly, every pattern that most significantly distinguishes among the clinical statuses involves CD68 macrophages (“CD68 macs”) (Fig. 6E). The SPACE covariation plot reveals that CD68 macs tend to cluster together in relatively small regions of each FoV and that regions with a high abundance of CD68 macs are enriched in postmortem and resection samples compared to biopsy material (Fig. 6F). We confirmed this by comparing the observed fraction of neighborhoods with ≥ 90% CD68 macs against null expectations for every sample (Fig. 6G).

We then investigated the ensemble that most significantly distinguishes the clinical statuses: neutrophil expression of IDO1, CD68 mac abundance, and CD11b/c CD206 macrophage (triple-positive “TP mac”) expression of PD-L1 (Fig. 6E). The covariation plot reveals that expression of PD-L1 by TP macs in the presence of CD68 macs is enriched in postmortem and resection samples vs. biopsy, while TP mac expression of PD-L1 in the absence of CD68 macs is depleted (Fig. 6H). We confirmed this by explicitly modeling TP mac expression of PD-L1 as a function of local CD68 mac abundance and clinical status. In postmortem and resection samples, PD-L1 on TP macs increases with local CD68 mac abundance, while the opposite is true in biopsy samples (slopes significantly differ, P = 0.0003, Fig. 6I). We also confirmed this visually. The highest PD-L1 expressing TP macs occur near CD68 macs in postmortem and resection samples but away from CD68 macs in biopsy samples (Fig. 6J, white arrows). A similar pattern involving CD68 macs and IDO1 expression on neutrophils was also suggested by SPACE (Fig. 6H) and confirmed by linear modeling (P = 1.73 × 10^−9^). Because such custom modeling requires a pre-existing hypothesis, nuanced patterns that escape the scope of existing spatial analysis platforms would previously have been nearly impossible to detect. However, SPACE can detect quite complex and subtle spatial patterns that correlate with clinical status.

## Discussion

High-content spatial representations of biological tissues contain enormous amounts of information on tissue architecture and cellular arrangement. To our knowledge, SPACE is the first analysis platform able to quantify spatial patterns of any number of participants and any degree of complexity, including context-dependent associations and quantitative gradients and orientations. These patterns may represent deviation from random assortment in a single specimen using cisMI, or distinctions among sample groups of specimens using transMI. In either case, SPACE ranks the exhaustively searched patterns, narrowing the enormous amount of information in the original data to a short list of the most significant patterns, which can then be studied in detail individually using the covariation plot.

SPACE is compatible with many input data types and pre-processing steps. In this paper alone, we utilize multiplex imaging in both 2d and 3d, via fluorescence- and heavy metal isotope-based techniques, as well as spatial transcriptomics. Segmented cells, 3d surface renderings, and raw biomolecule quantification can all be analyzed, in full image or condensed table formats. Multiple representations of the same specimens can be analyzed simultaneously to combine their unique advantages.

SPACE discovers complex patterns that are not accessible to existing analytical techniques. For example, in the mouse LN, we show that pairwise associations cannot describe the context-dependent nature of the opposing CD4 vs. CD8 T cell gradient in the paracortex nor the consistent DZ vs. LZ orientation of GCs with respect to the capsule, because both patterns are relative to a third cell type. Nor can MEs describe these patterns, because both patterns involve continuous features that are not easily divided into discrete regions. Again, in human intestinal cancer, the tradeoff between IGHG4 and SFRP2 transcripts specifically where both are abundant is not detected by pairwise correlation nor by ME delineation. Thus, SPACE is not an incremental improvement in accuracy or power for existing analyses – it is a new analysis approach that detects novel types of spatial patterns altogether.

Therefore, even in the re-analysis of existing data sets, SPACE may provide additional insights of biological importance. In a mouse tumor, the biased association of CD4 T cells toward blood vessels recalls prior literature in which interactions between different T cell subsets and vasculature dictate trajectories of tumor growth [41] or infection [42]. In the human intestinal cancer specimen, the broad colocalization but local anticorrelation of IGHG4 and SFRP2 may reflect the involvement of humoral immunity in combating the cancer. Downregulation of SFRP2 expression promotes colorectal cancer cell proliferation [44–45]. Meanwhile, cells expressing immunoglobulins (involving IGHG4), such as B cells and IgG plasma cells, infiltrate and help control colorectal tumors [46–47]. Such targeting of B cells and IgG plasma cells to regions of cancerous proliferation would lead to an abundance of IGHG4 where SFRP2 is expressed broadly, but especially where SFRP2 is repressed locally.

The most insightful spatial patterns may not be those that differ from random assortment, but rather those that differ systematically among sample groups. SPACE is one of the first spatial analysis platforms to make such statistical comparisons directly, as we demonstrate using the human TB data set. SPACE finds that local accumulation of CD68 macs is particularly prevalent in the more severe disease cases (therapeutic resections and postmortem samples) as opposed to the less severe (diagnostic biopsies), which has also been observed in non-human primates [48]. Near these CD68 macs, immunoregulatory molecule expression, including PD-L1 on TP macs and IDO1 on neutrophils, increases in more severe disease cases but decreases in less severe cases. This suggests that CD68 macs may play a crucial and bifurcating role in bacterial containment: CD68 macs switching from a pro-inflammatory to a pro-regulatory role, perhaps as the result of some density-driven feedback, may cause disease to progress. This is accompanied by a loss of beta then alpha diversity, suggesting that as immunological control deteriorates, ME integrity is lost as immune cell types first intermix and then begin to disappear from the granuloma altogether.

Speculative interpretations of the patterns discovered in this study should be tempered, however. Sample sizes are small – just one for the cisMI demonstrations, and just 15 patients for the TB data. Potentially crucial metadata on co-morbidities, ongoing treatments, etc. are missing. Even without these shortcomings, static patterns are rarely sufficient evidence of dynamic processes. Thus, further work would be required to test the hypotheses proposed here. Nonetheless, SPACE presents a substantial advance in the ability to search for complex spatial patterns quickly and exhaustively, streamlining the process of hypothesis generation.

Although we developed SPACE with high-content tissue images in mind, SPACE can census and analyze any image-based data, from super-resolution microscopy of single cells to remote sensing of earth’s surface. In fact, the information-theoretic calculation of cis- and transMI is agnostic to the source of the data, whether spatial or not. So long as randomized null expectations can be credibly simulated, SPACE essentially performs a non-parametric test for divergences between distributions. Whereas common non-parametric procedures only compare the first moment of 1-dimensional distributions, SPACE compares the full shape of multi-dimensional distributions. Thus, SPACE could prove to be a highly useful statistical innovation. SPACE is available as a free, open-source R package that can be widely adopted and modified to support advanced analytical capabilities in a variety of scientific fields.

## Materials and Methods

### SPACE Computing Environment and Availability

SPACE is an R package, written in R version 4.3.1 and developed on a laptop with 16GB RAM and an 11th Gen Intel® Core™ i9-11900H 2.50GHz processer. SPACE is open source and freely available from [https://github.com/eschrom/SPACE], along with a tutorial demonstrating the usage of each function.

### Details of SPACE Censusing

Input images of biomolecular expression must be in 8-bit grayscale .tif format, using multi-channel stacks. Input images of color-coded objects must be in RGB .tif format. Tables of centroids must be in .csv format.

Before censusing, the size and number of neighborhoods to draw are chosen. Given the image resolution in µm/pixel (which may differ in the X, Y, and Z dimensions), neighborhood size is chosen to reflect a specified radius in µm. (For applications outside the microscopic scale, a different default unit of length may be chosen.) A range of radii may be used to draw neighborhoods of varying sizes. For each neighborhood size, the number of neighborhoods to draw is chosen based on coverage: the expected number of neighborhoods that include each non-background pixel. Higher coverage results in more neighborhoods and higher statistical power but longer computational times. An upper limit of 5x coverage is recommended, above which pseudo-replication becomes a serious statistical flaw.

To census a specimen, seed points are chosen as the centers of each neighborhood. Seed points may be distributed evenly across all non-background areas or targeted to specific objects or biomolecules. Within these constraints, seed points are distributed randomly, such that neighborhoods may overlap but no two are identical.

The amount of each variable in each neighborhood is summarized in the census. Additionally, contiguous patches of each variable in each neighborhood, respecting truncation by neighborhood boundaries, are recorded in a patch list. Randomized censuses are simulated by reassorting the individual patches across the neighborhoods at random, while holding the total non-background content of each neighborhood fixed.

When censusing, biomolecules may be considered independent or linked. Independent biomolecules are considered single variables whose overall expression may be related spatially to other variables. For each independent biomolecule, a binarization threshold for its expression is first calculated across the full image via the IsoData algorithm. This threshold is used to delineate patches of the biomolecule within each neighborhood, but the true quantitative expression intensity within each patch is recorded in the patch list.

Linked biomolecules are considered separate variables when expressed on separate objects. For example, “Biomolecule A on Object X” is considered a separate variable from “Biomolecule A on Object Y.” Patches of linked biomolecules are defined by the patches of their corresponding objects. A table in .csv format can be provided to specify which biomolecule-object combinations are of interest to measure and record in the census.

### Details of CisMI Measurement

For a given ensemble, the distribution of co-occurrence is extracted from the census, along with distributions for all subsets of the ensemble. For example, for the ensemble of Variables A, B, and C, distributions are constructed for ABC, AB, BC, AC, A, B, and C. The entropy of each distribution (denoted H(ABC), H(AB), etc.) is calculated. Entropies are added and subtracted according to the definition of interaction information (denoted I(ABC)). In the three-variable case, interaction information is given by:

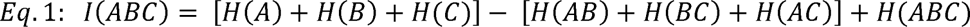

Accurate quantification of interaction information requires accurate quantification of entropy. SPACE uses the “histogram estimate” of entropy, for which the abundance of each variable in each neighborhood must be rounded into discrete bins. Bins have constant widths on the interval from 0 – 100%, where 100% is the maximum abundance of a given variable across all neighborhoods. For example, if there are 5 bins per variable, then a variable’s abundance in each neighborhood is rounded into 0-20%, 20-40%, 40-60%, 60-80%, or 80-100% of its maximum abundance across all neighborhoods. For a constant number of bins per variable b, the total number of bins B in a distribution depends on the number of variables that distribution has. For b = 5, a 1-variable distribution has B = 5 bins, a 2-variable distribution has B = 5^2^ = 25 bins, a 3-variable distribution has B = 5^3^ = 125 bins, and so on.

For the histogram estimate of entropy to be accurate, the number of neighborhoods (i.e., observations) N must greatly exceed the number of total bins B. The B/N ratio will be more constraining for ensembles that include more variables. To ensure that entropy is estimated accurately for all ensembles, even those including the maximal number of variables D, b is chosen so that B is always less than or equal to one-tenth N:

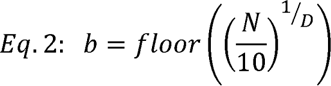

Even with a well-chosen number of bins per variable, the histogram estimate of entropy is still slightly biased downward. A variety of correction factors have been used historically to counteract this bias, such as adding a term based on the B/N ratio [49]. SPACE adds the following term to all entropy estimates to finalize the entropy calculation:

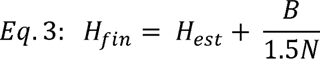

With accurate entropy estimates, interaction information can also be estimated accurately. However, factors other than non-random spatial patterning can create interaction information, e.g., cellular or structural size and morphology, tissue geometry, and the compositional nature of objects in space. To control for these factors, interaction information is also calculated for many randomized censuses, where these factors remain but spatial patterning is removed. As few as 10 randomized censuses may suffice for consistent results, but ≥ 100 is usually recommended when feasible. Subtracting the mean randomized interaction information from the observed interaction information therefore isolates the interaction information due to spatial patterning specifically. We call this measure cisMI.

The statistical significance of an ensemble’s cisMI is assessed by expressing the observed interaction information as a Z score relative to the distribution of randomized interaction information values. The probability of a value more extreme than this Z score is the ensemble’s P value. For Z scores that are too large to measure a P value from the Gaussian cumulative density function computationally, quadratic regression is used to estimate the log P value. When cisMI is measured for many ensembles, P values are corrected for multiple testing following the Benjamini-Hochberg procedure.

### Details of TransMI Measurement

For a given ensemble, the distribution of co-occurrence is extracted from the censuses of each specimen. For each pair of specimens, the difference between their co-occurrence distributions is quantified via KL divergence. Like entropy, KL divergence requires the abundance of each variable in each neighborhood to be rounded into discrete bins. The number of bins per variable is chosen exactly as for cisMI.

KL divergence also requires that any bin with a nonzero probability in one distribution also has a nonzero probability in the other. Regularization guarantee that this condition is met. For each distribution, the empirical counts of neighborhoods in each bin are used to calculate the Chao-Jost coverage estimator C [50], and the empirical bin probabilities are normalized to C. The remaining 1-C probability mass is distributed uniformly across all possible but unobserved bins. KL divergence is then calculated for these regularized distributions. Because KL divergence is not symmetric, it is calculated in both directions and averaged. We call this measure transMI.

Once transMI has been measured for every pair of specimens, a fully connected network is constructed. The specimens are the vertices, and the undirected edges E between specimens i and j out of the full set of specimens S are weighted according to:

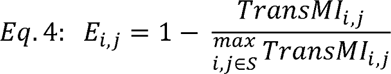

As such, pairs of specimens with dissimilar spatial patterning and therefore high transMI are assigned low edge weights, and pairs of specimens with similar spatial patterning and therefore low transMI are assigned high edge weights. Because it is assumed that the specimens belong to two or more discrete sample groups, the modularity of this network is then calculated. High modularity indicates that the ensemble’s spatial patterning differs systematically across sample groups. Because high modularity can arise by chance, the edge weights are permuted to yield randomized estimates of transMI modularity. Typically, 100 randomized estimates are used.

The statistical significance of an ensemble’s transMI modularity is assessed by expressing the observed transMI modularity as a Z score relative to the distribution of randomized transMI modularity values. The probability of a value more extreme than this Z score is the ensemble’s P value. For Z scores that are too large to measure a P value from the Gaussian cumulative density function computationally, quadratic regression is used to estimate the log P value. When transMI is measured for many ensembles, P values are corrected for multiple testing following the Benjamini-Hochberg procedure.

### Details of Pattern Characterization

To describe the pattern formed by a particular ensemble, the ensemble’s distribution of co-occurrence is generated, for both focal and reference censuses. For cisMI, the observed census is focal and a randomized census is the reference. For transMI, the pooled censuses from one sample group are focal, and pooled censuses from another sample group are the reference. A curve is then drawn on this distribution through the areas of highest focal neighborhood density, using a self-organizing map. To enforce that this curve is a 1-dimensional manifold in D-dimensional space (where D is the number of variables in the ensemble), the self-organizing map nodes are arranged in an N x 1 grid. By default, N is 1000, this grid is non-toroidal, and 50 iterations of the self-organizing map are run with a learning rate decreasing from 0.5 to 0.1.

The curve that results from the self-organizing map is really a sequence of N nodes described by coordinates in D-dimensions, where each dimension represents the abundance of one of the variables in the ensemble. Each focal neighborhood is assigned to its closest node as measured by Euclidean distance. Focal neighborhoods are ordered according to their closest node, with ties broken at random. The rolling mean and 95% confidence interval for the abundance of each variable is plotted in the neighborhood order determined by the self-organizing map (i.e., the latent path) to create the covariation plot.

Although reference neighborhoods do not affect the placement of the self-organizing map curve, each reference neighborhood is also assigned to its closest node as measured by Euclidean distance. For each node n, the I_n_ focal and J_n_ reference neighborhoods assigned to that node are weighted based on their individual distances d to the node, such that closer neighborhoods are weighted more heavily. Then, the log ratio of the weighted fractions of focal vs. reference neighborhoods assigned to the node are used as the node’s enrichment score E_n_:

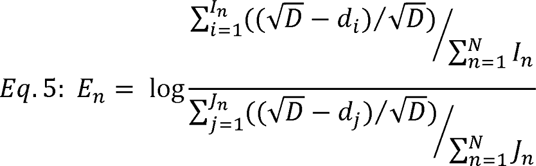

If a higher fraction of focal neighborhoods is clustered more closely around a node compared to reference neighborhoods, the enrichment score will be positive. If a lower fraction of focal neighborhoods is clustered more loosely around a node compared to reference neighborhoods, the enrichment score will be negative. The enrichment scores are ordered and repeated as necessary to follow the latent path. The rolling mean of the enrichment scores are then added to the covariation plot, to highlight features that are particularly different between the focal and the reference censuses. Because the self-organizing map is only drawn with respect to the focal census, swapping the focal and reference censuses may not yield an identical covariation plot nor a precisely inverted enrichment plot.

To highlight the regions of the image that correspond to certain features of the covariation plot, these features are bracketed by manually chosen breakpoints along the latent path. This defines a partitioning of the neighborhoods of the focal census. These neighborhoods are then traced back to their original locations in their original specimen’s image, enabling a voting process at the pixel level which assigns each pixel into one of the regions (or MEs) defined by the breakpoints along the latent path. Pixels are then recolored by the ME to which they belong, revealing a new image of ME organization.

### Mouse Lymph Node Cellular Phenotyping

The 42-parameter image of the mouse popliteal lymph node is publicly available [https://zenodo.org/records/10870403]. The JOJO nuclear stain and CD45 membrane stain were supplied to Cellpose 2.0 [37] to obtain a segmentation mask of individual cells. Using an ImageJ macro, the mean fluorescent intensity (MFI) was calculated for each single cell, for 20 of the 42 original fluorescent markers. A .csv table of lineage expectations was manually created, in which the columns represent these 20 markers, and the rows represent 18 known cell types that are identifiable using these 20 markers. For each entry, 1 indicates that the cell type is expected to express the marker, 0 indicates that the cell type is not expected to express the marker, and NA indicates that the cell type may or may not express the marker.

Using an R script, a binarization threshold was calculated using the MFI distribution across single cells for each marker, via the IsoData algorithm. Across all cells and markers, the probability that cell i expresses marker j P_i,j_ was calculated using cell i’s expression of marker j MFI_i,j_ and the binarization threshold for marker j T_j_ according to a sigmoid curve as follows:

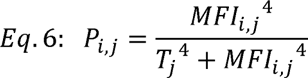

By comparing each cell’s expression probabilities to the manually defined lineage expectations, each cell was assigned a probability score of belonging to each of the 18 types. The probability score that cell i belongs to type k P_i,k_ was calculated from cell i’s expression probabilities P_i,j_ and cell type k’s expected expression profile L_k,j_ from the lineage expectation table as follows:

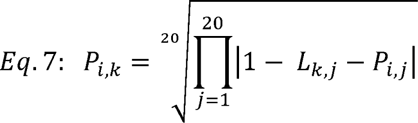

For each instance where L_k,j_ was NA, the corresponding term was omitted from the product, and the index of the radical was reduced by 1.

Once P_i,k_ was calculated for all cells and types, each cell was assigned to the type for which it had the highest probability score, if that score exceeded 0.5. Otherwise, the cell was assigned to the “unknown” 19^th^ type. In principle, unknown cells can be subject to unsupervised clustering approaches to define more cell types not included in the original lineage expectations. Here, however, unknown cells were left as such.

### Mouse Lymph Node SPACE Analysis

On the cellular segmentation mask color-coded by the 19 cell types, 8609 neighborhoods of radius 10 µm were drawn using the SPACE ‘census_image’ function. The radius of 10 µm was chosen to capture close cellular contacts, since the lymphocytes that comprise most of the LN are 5-7 µm in diameter. The number of neighborhoods was chosen to achieve 5x tissue coverage (i.e., each non-background pixel is included in an average of 5 neighborhoods), so that the tissue is thoroughly covered without excessive pseudo-replication.

From this census and its corresponding patch list, cisMI was measured for all ensembles of one, two, or three cell types using the SPACE ‘measure_cisMI’ function. This function was also used to measure cisMI for the specific 4-element ensemble of CD4+ T cells, CD8+ T cells, cDC1s, and cDC2s. Ensembles for further exploration were chosen based on P value and/or cisMI magnitude. Covariation plots were generated using the SPACE ‘learn_pattern’ function. Visualizations of representative regions were generated by aligning fluorescent staining with the segmented objects and then zooming and cropping in ImageJ.

Censusing and cisMI measurement was performed identically using a table of cell centroids. To compare the table-based and image-based results, the cisMI Z scores for all 1159 ensembles of one, two, or three variables were compared via linear regression.

### Mouse Lymph Node SPACE Benchmarking

To perform a generic analysis representative of existing pairwise approaches, the number of neighbors of each cell type were counted within 10 µm of each cell. For each cell type, this gave an average number of neighbors of each cell type. The cell types were then permuted among the fixed centroids and this measure was repeated 100 times. For each cell type pair, the true average number of neighbors was expressed as a Z-score of the 100 null measurements.

To perform a generic analysis representative of existing ME approaches, the table-based neighborhoods of each cell were subject to 50 K-means clustering runs at each k value from 2 to 25, where clusters are synonymous with MEs. Each iteration, the cell type compositions of each ME were used to count the number of key features representative of the patterns identified by SPACE (Table S3), as well as the Silhouette score.

### Mouse Tumor 3D SPACE Analysis

A volumetric image of the mouse MC38 tumor was collected using the Ce3D protocol [Li et al. 2017] for 6 fluorescent markers: MHCII, Lyve1, CD3, CD4, CD8, and CD31. After cropping the full image to a smaller representative region in Imaris 10.0, the CD31 image was masked on MHCII and Lyve1 to remove spillover from these channels. The Imaris surface module was used to create 3d renderings of the CD31 blood vessels and the CD3 T cells. The CD3 T cells were then partitioned into CD4 T cells (CD4 MFI > 180 and CD8 MFI < 30) and CD8 T cells (CD4 MFI < 180 and CD8 MFI > 30). The ratio of total T cell volume to total blood vessel volume (79%) was used to estimate the number of expected vascular endothelial cells (3464) relative to the number of T cells (4388). The Imaris spots module was used to distribute 3464 spots along the CD31+ blood vessel surfaces. The XYZ coordinates of each T cell and blood vessel spot were then exported from Imaris.

These coordinates were loaded into SPACE, and 1569 neighborhoods of radius 50 µm were drawn using the SPACE ‘census_table’ function. The radius of 50 µm was chosen to capture broad cellular associations. The number of neighborhoods was chosen to achieve 5x tissue coverage (i.e. each object is included in an average of 5 neighborhoods) so that the tissue volume is covered thoroughly without excessive pseudo-replication.

From this census and its corresponding patch list, cisMI was measured for the ensemble of all three object types (CD4 T cells, CD8 T cells, and blood vessels) using the SPACE ‘measure_cisMI’ function. The covariation plot for the three-variable ensemble was created using the SPACE ‘learn_pattern’ function.

To confirm the patterns found by SPACE, T cells were separated into those < 50 µm from the nearest blood vessel vs. those > 50 µm. The number of CD4 vs. CD8 T cells in each class was compared using a linear model with a binomial likelihood. The distance to the nearest vessel for CD4 vs. CD8 T cells in each class was compared using a linear model with a Gaussian likelihood.

### Visium Human Intestinal Colorectal Cancer Analysis

The 6.5mm x 6.5mm section of human colorectal cancer was publicly available from 10x Genomics [https://www.10xgenomics.com/datasets/human-intestine-cancer-1-standard], along with Moran’s I for individual transcripts, K-means clusterings, and differential abundance analyses across the 2660 spots. After filtering for transcripts that appear in at least half of the spots, 6914 transcripts remained. Using each spot as a seed point, 2660 neighborhoods of radius 300 µm were drawn using the SPACE ‘census_table’ function. The large neighborhood size was chosen because the target comparison metric, Moran’s I, is a global measure. CisMI was calculated for each transcript individually using the ‘measure_cisMI’ function. After filtering out 36 additional transcripts with negative Moran’s I, the Moran’s I values were log-transformed and compared with cisMI via standardization and linear regression.

The publicly available differential abundance analysis was used to identify 13 transcripts that significantly differ from their global means in at least 4 of the 8 MEs. Using each spot as a seed point, 2660 neighborhoods of radius 150 µm were drawn using the SPACE ‘census_table’ function. The smaller neighborhood size was chosen because the target comparison metric, k-means MEs, were drawn at the scale of individual spots. CisMI was calculated for ensembles of up to three transcripts using the ‘measure_cisMI’ function. The covariation plots for specific ensembles were drawn using the ‘learn_pattern’ function.

After drawing gates on the covariation plots to assign regions of interest, these regions were mapped back onto the tissue by assigning each spot to its most similar self-organizing map node. After counting the number of mapped spots belonging to each ME, a 2×2 table reported the total number of spots that belong to the gated covariation region, the top-matching ME, both, or neither. Cohen’s κ was calculated from this table to quantify the agreement between the gated feature(s) of the covariation plot and the top-matching ME.

Correlations between pairs of transcripts were measured across all 2660 spots by standardizing the transcript counts and performing linear regression.

### Human TB Analysis

The 30 TB FoVs, including phenotyped cells, ME demarcation, and functional marker expression, were publicly available from [McCaffrey et al 2022]. The cellular alpha and beta diversity of each FoV were measured using the ‘alpha_diversity’ and ‘beta_diversity’ SPACE functions. Diversity metrics were compared across clinical statuses using a linear model with clinical status as a categorical predictor.

On each FoV, 1825 neighborhoods of radius 10 µm were drawn using the SPACE ‘census_image’ function. The radius of 10 µm was chosen to capture close cellular associations. The number of neighborhoods was chosen to achieve 5x tissue coverage (i.e. each object is included in an average of 5 neighborhoods) so that the tissue volume is covered thoroughly without excessive pseudo-replication.

The total list of cell types, MEs, and cell-type-specific marker expression was reduced to include variables that occur in at least 3 of the 6 resection FoVs, at least 3 of the 6 postmortem FoVs, and at least 9 of the 18 biopsy FoVs. TransMI was measured for every ensemble of up to three of these variables at a time using the SPACE ‘measure_transMI’ function. Covariation plots were created using the SPACE ‘learn_pattern’ function.

To measure the frequency of CD68 mac aggregates in each FoV, the fraction of neighborhoods with ≥ 90% CD68 macs from the observed census was divided by the same fraction from a randomized census, generated using the SPACE ‘plot_dist’ function. The logarithms of these ratios were compared across clinical statuses using a linear model with clinical status as a categorical predictor.

To quantify how cell-type-specific marker expression (PD-L1 on TP macs or IDO on neutrophils) depends on local CD68 mac abundance and/or clinical status, all neighborhoods with non-zero cell-type-specific marker expression were pooled from the observed censuses. The log of cell-type-specific marker expression was then regressed on the interaction of quantitative local CD68 mac abundance and categorical clinical status (where resection and postmortem were pooled), with a random effect of patient ID.

## Acknowledgments

This research was supported by the Division of Intramural Research of NIAID, NIH, the Center for Cancer Research, NCI, NIH, and the Chan-Zuckerberg Initiative.

## Author Contributions

ECS conceived of the work, developed the mathematical approach, wrote the computer code, analyzed the primary data, interpreted the data analysis, and wrote the manuscript.

EFM contributed primary data, interpreted the data analysis, and edited the manuscript.

VS wrote the computer code.

AJR contributed primary data, interpreted the data analysis, and edited the manuscript.

HI contributed primary data and interpreted the data analysis.

AAM contributed primary data.

ES interpreted the data analysis and edited the manuscript.

LA interpreted the data analysis and edited the manuscript.

NT analyzed the primary data.

SG analyzed the primary data.

RNG conceived of the work, interpreted the data analysis, and edited the manuscript.

## Supplementary Figure and Table Legends

**Table S1.** Lineage marker expression expectations used for semi-supervised cellular phenotyping in the mouse LN. 1 indicates expected expression, 0 indicates expected lack of expression, and NA indicates no expectation.

**Table S2.** Centroid table of cellular positioning and phenotype in the mouse LN.

**Table S3.** Key features of ME compositions required to demonstrate patterns of the mouse LN organization discovered by SPACE.

**Figure S1.**
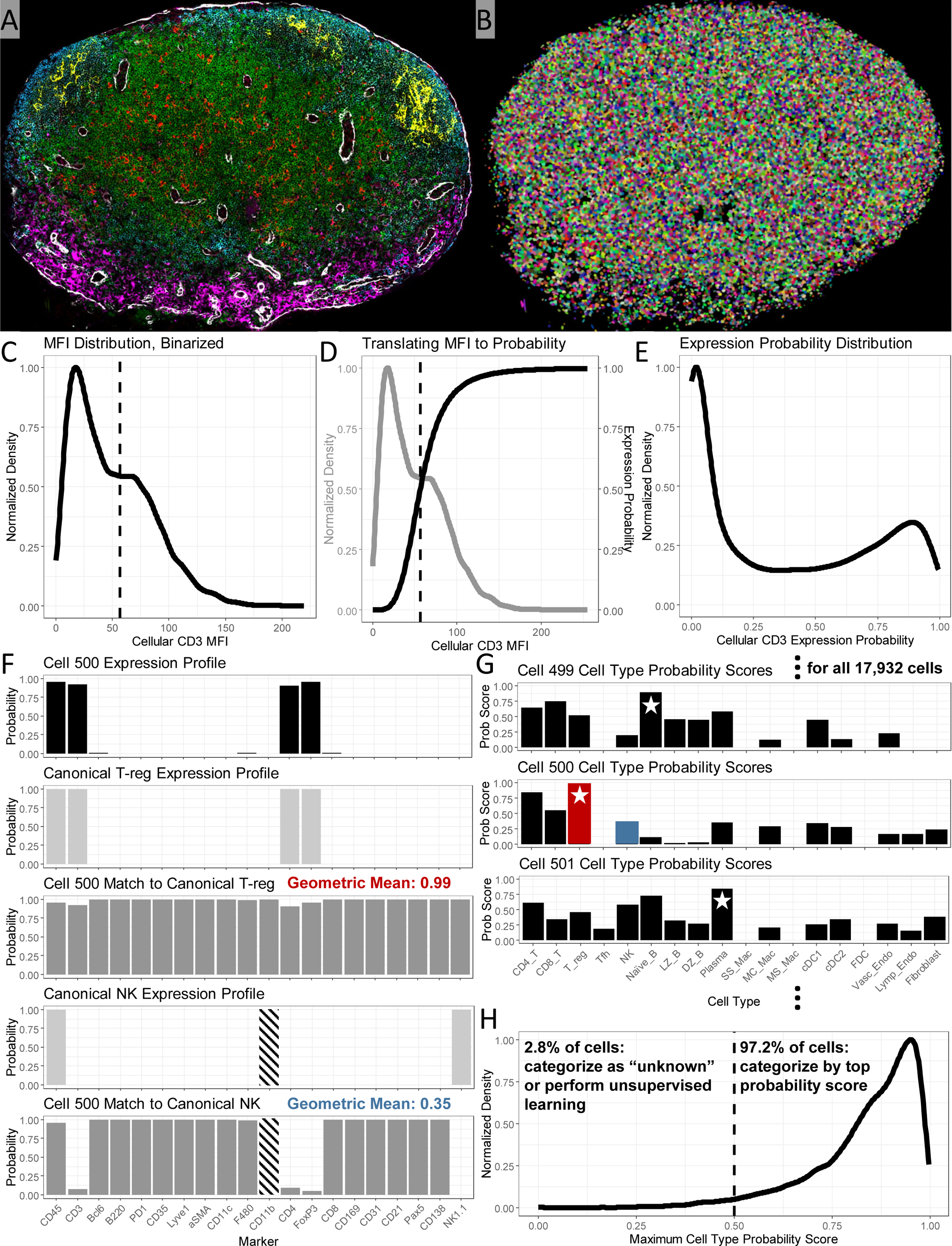
Cellular segmentation and semi-supervised phenotyping for the mouse LN. A) The original 42-plex IBEX image of a mouse popliteal LN. Six channels are shown: CD3 (green), B220 (blue), F480 (pink), DEC205 (red), CD21 (yellow), αSMA (white). B) Segmentation mask of single cells, generated in Cellpose 2.0 on default settings with JOJO as nucleus and CD45 as membrane. C) For each of 20 lineage markers in the reduced panel, the mean fluorescence intensity (MFI) across cells forms a distribution, for which a binarization threshold is chosen using the IsoData algorithm. CD3 is shown as an example. D) MFI is transformed into a probability-of-expression using a Hill function sigmoid curve with EC50 equal to the marker’s binarization threshold and Hill exponent equal to 4. E) The distribution of MFI across cells is transformed into a distribution of probability-of-expression, in which positive and negative cells are more clearly separated. F) For each cell, the expression probability is calculated for each marker. Cell 500 out of 17,932 is shown as an example. This expression profile is compared to the canonical profile for each expected cell type. T-reg and NK are shown as examples. In each comparison, the focal cell’s expression probabilities are unchanged for markers expected to be positive and subtracted from 1 for markers expected to be negative. The geometric mean of these values gives a probability score describing how likely the focal cell is to belong to the canonical type. Expression probabilities are omitted from the geometric mean when they are uninformative; for example, NK expression of CD11b is variable, so CD11b is uninformative when evaluating whether a cell belongs to the NK type. G) For each cell in the data set, a probability score is calculated for each canonical cell type, and the maximum score indicates the most likely type. H) The maximum score is close to 1 for most cells, indicating a good match. 97.2% of cells have a maximum score > 0.5 and are categorized into one of the canonical types. 2.8% of cells have a maximum score ≤ 0.5. Here, such cells are considered “unknown;’ however, unsupervised clustering approaches can be applied to this subset of cells to discover unexpected types.

**Figure S2.**
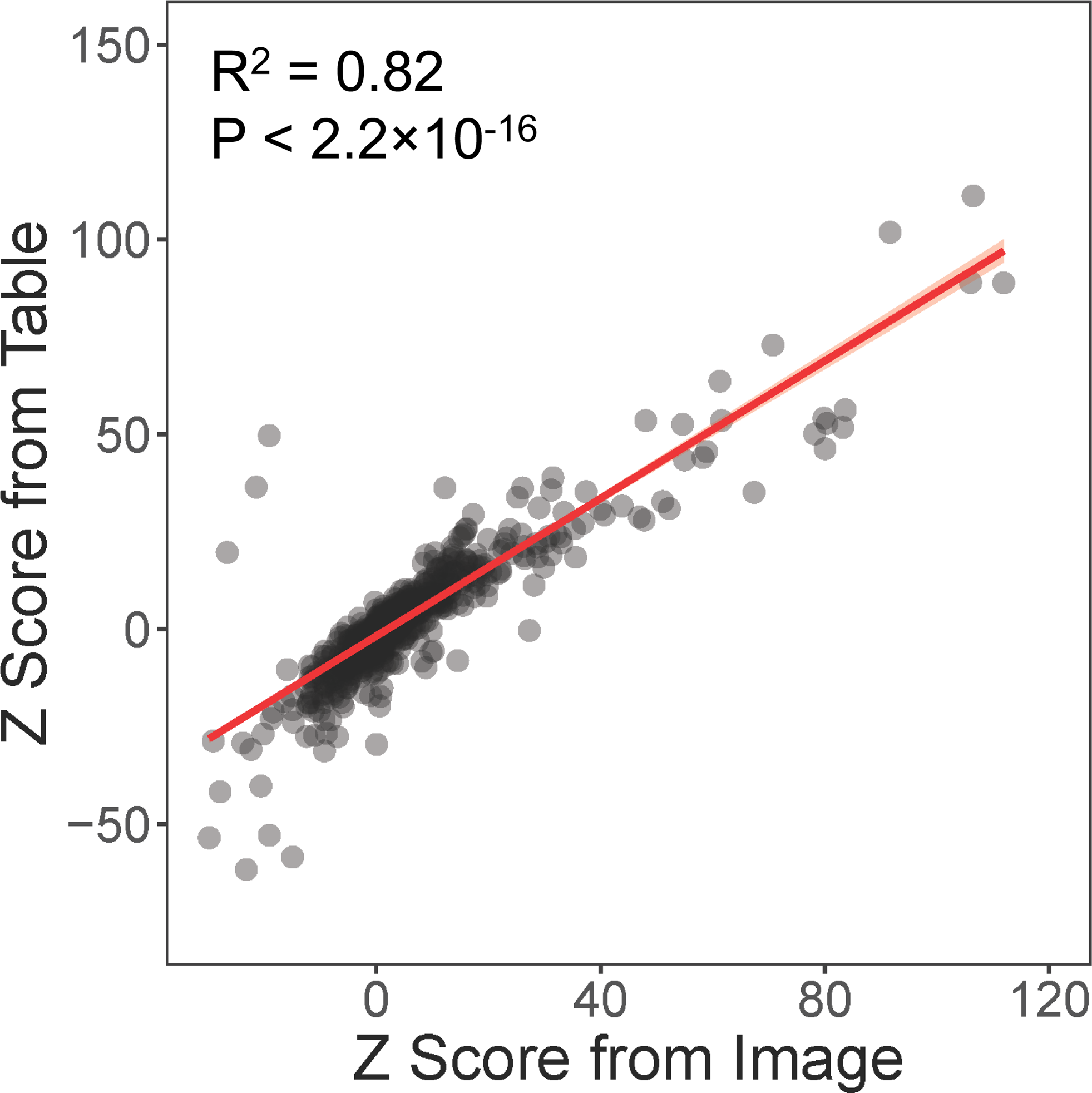
Comparison of cisMI Z scores measured by SPACE for all ensembles of up to 3 out of the 19 cell types in the mouse LN from the segmentation image vs. the centroid table, using a Gaussian linear model.

**Figure S3.**
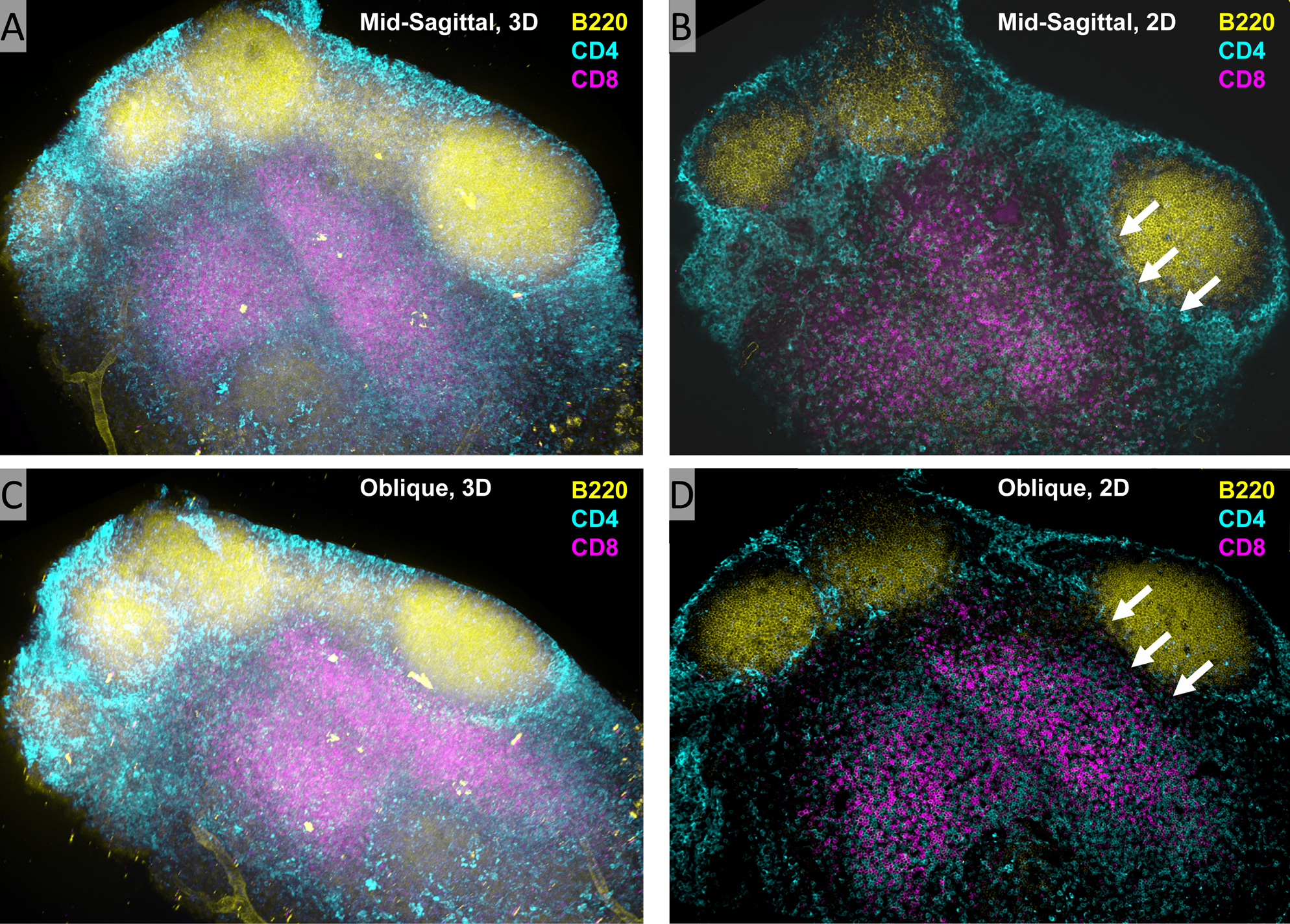
Volumetric image of a mouse LN to visualize different slicing angles. A) The full 3d volume viewed from a mid-sagittal angle shows the inside-out gradient of CD8 to CD4 T cells. B) A single 2d slice viewed from a mid-sagittal angle also shows the inside-out gradient of CD8 to CD4 T cells. C) The full 3d volume viewed slightly oblique to a mid-sagittal angle shows a right-to-left gradient of CD8 to CD4 T cells. D) A single 2d slice slightly oblique to a mid-sagittal angle also shows a right-to-left gradient of CD8 to CD4 T cells. White arrow show where the CD4 T cells that separate the CD8 T cells from the B follicle disappear fully on the right, but not fully on the left.

**Figure S4.**
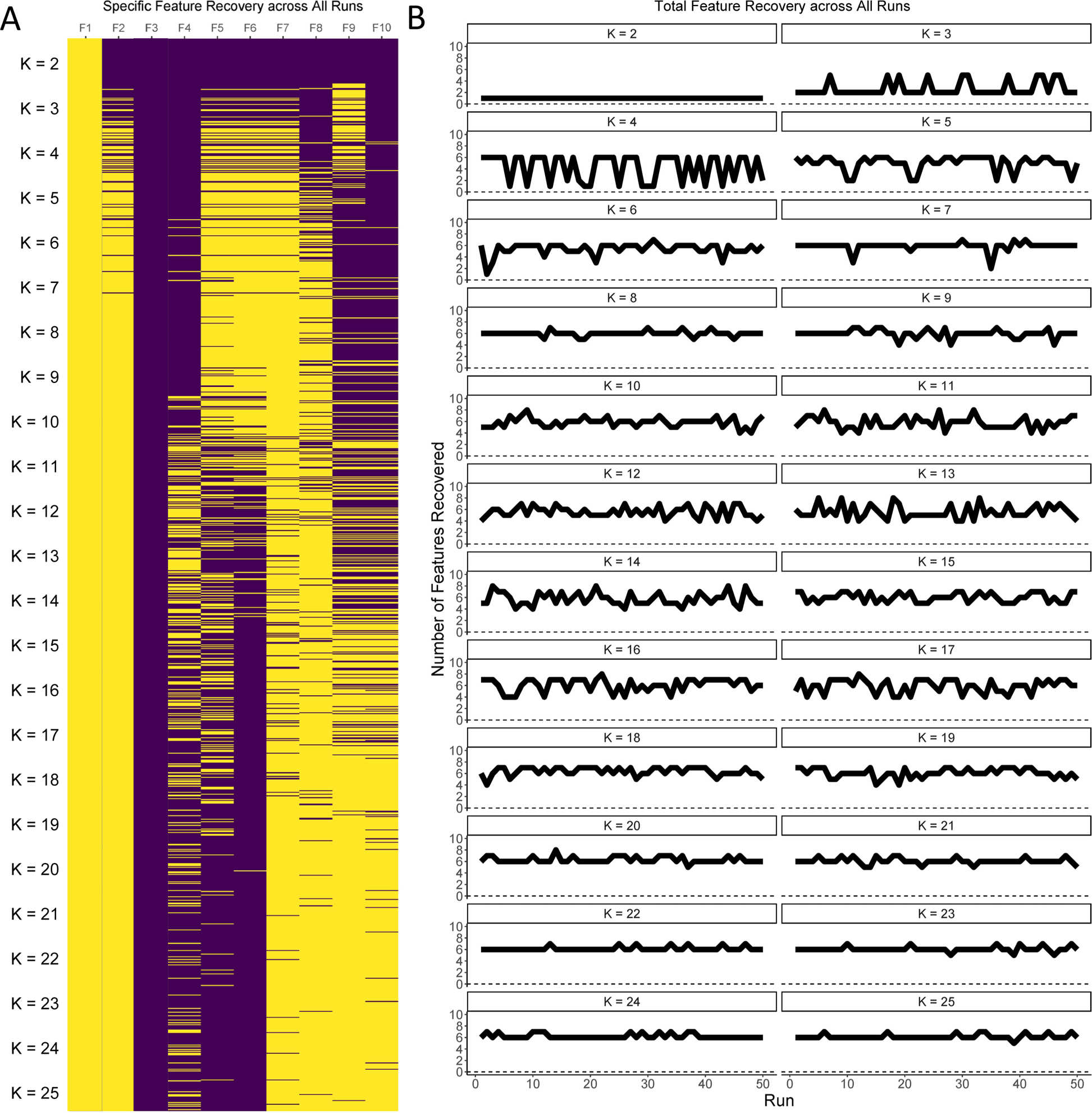
All K-means clustering runs to define MEs on the mouse LN across K values from 2 to 25. A) Specific key features recovered by each clustering run. B) Total key features recovered by each clustering run.

**Figure S5.**
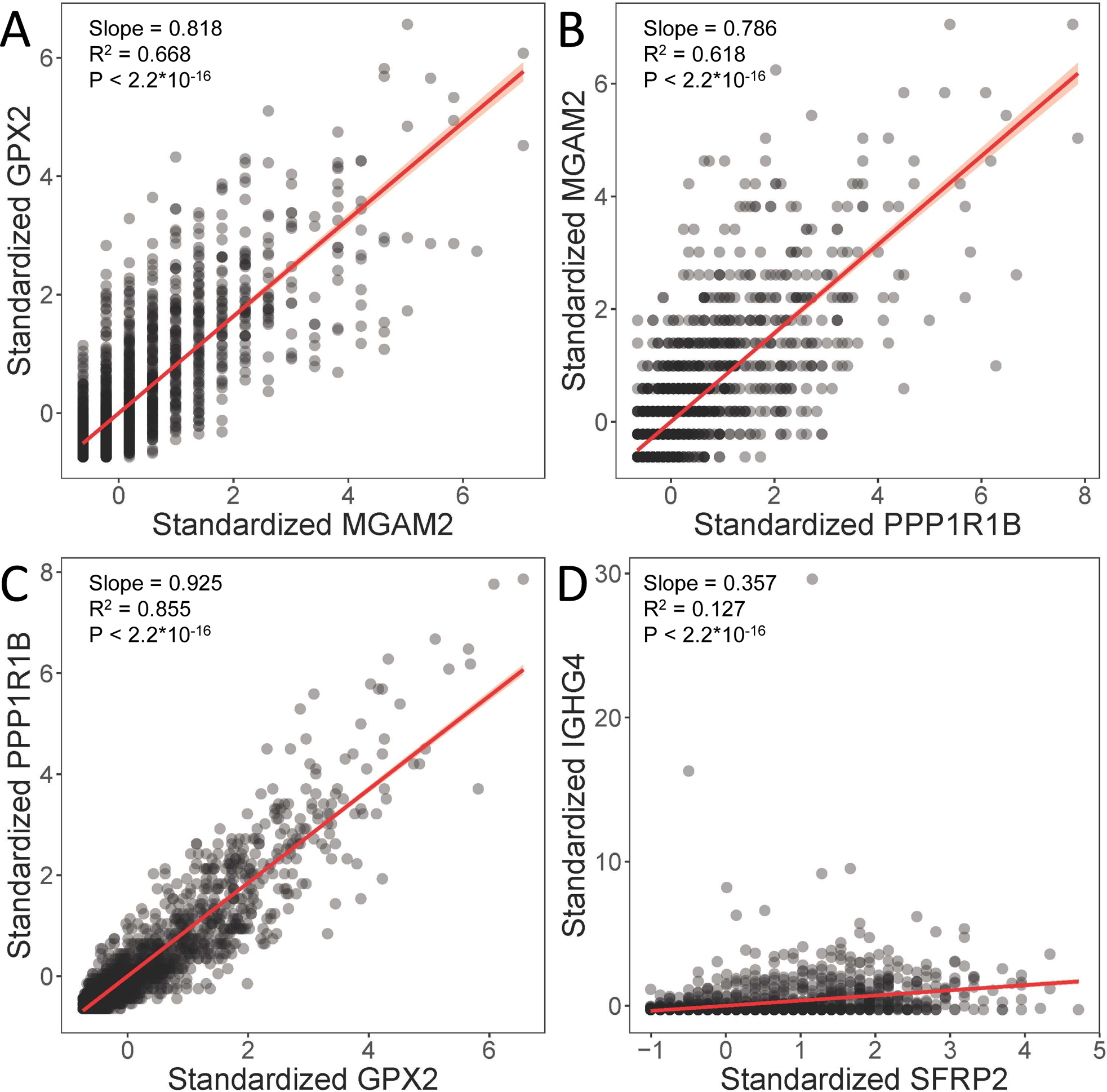
Pairwise correlations of transcript abundance across the 2660 spatial spots in the human intestinal cancer sample. A) Comparison of standardized counts of GPX2 and MGAM2. B) Comparison of standardized counts of MGAM2 and PPP1R1B. C) Comparison of standardized counts of PPP1R1B and GPX2. D) Comparison of standardized counts of IGHG4 and SFRP2. All comparisons were made with a Gaussian linear model.

